# Loss of TMEM55B modulates lipid metabolism through dysregulated lipophagy and mitochondrial function

**DOI:** 10.1101/2024.11.12.623285

**Authors:** Yuanyuan Qin, Sheila Teker, Nilsa La Cunza, Elizabeth Theusch, Neil Yang, Leela Venkatesan, Julia Su, Ronald M. Krauss, Aparna Lakkaraju, Aras N. Mattis, Marisa W. Medina

## Abstract

Lipophagy is a form of selective autophagy that targets the lipid droplets for lysosomal decay and has been implicated in the onset and progression of metabolic dysfunction-associated steatotic liver disease (MASLD). Factors that augment lipophagy have been identified as targets for MASLD therapeutic development. TMEM55B is a key regulator of lysosomal positioning which is critical for the lysosome fusion with the autophagosome, but less studied. Here, we demonstrate that inhibition of TMEM55B in murine models accelerates MASLD onset and progression. In cellular models, TMEM55B deficiency enhances lipophagy, leading to increased fatty acid release from lysosomes to mitochondria but simultaneously impairs mitophagy, causing an accumulation of dysfunctional mitochondria. This imbalance leads to increased lipid accumulation and oxidative stress, worsening MASLD. These findings underscore the importance of lysosomal positioning in lipid metabolism and suggest that augmenting lipophagy may exacerbate disease in the context of mitochondrial dysfunction.

## INTRODUCTION

Lipophagy is a form of selective autophagy that specifically targets lipid droplets to maintain cellular lipid homeostasis. There are two forms of lipophagy: macrolipophagy in which lipid droplets are engulfed by autophagosomes after which they fuse with lysosomes^1^, and microlipophagy, where triglycerides are directly extruded from lipid droplets into lysosomes^2^. Once inside the lysosome (or autophagolysosome), lysosomal enzymes break down lipid droplet-derived triglycerides into fatty acids, which may be directed to the mitochondria for energy production or used in other cellular processes^1^. By breaking down lipid droplets, lipophagy also prevents the accumulation of toxic lipid species, particularly important for mitigating lipotoxicity. Additionally, by targeting damaged or dysfunctional lipid droplets, lipophagy helps maintain lipid and energy homeostasis^3^. Based on these critical roles, impaired lipophagy has been implicated in the development and progression of metabolic dysfunction-associated steatotic liver disease (MASLD) in human patients and pre-clinical models^4–7^.

As the leading cause of chronic liver disease in the United States^8^, MASLD encompasses a spectrum of liver disorders initiated from excess hepatic lipids (i.e., hepatic steatosis), which can progress to metabolic-associated steatohepatitis (MASH), fibrosis, and cirrhosis^9^. The central basis of MASLD pathogenesis is the dysregulation of hepatic lipid metabolism^10^. Given the role of lipophagy in releasing triglycerides stored in lipid droplets, agents that elevate lipophagy have emerged as potential therapeutics for the treatment of MASLD^11^. Notably, the effectiveness of this mechanism requires functional mitochondria to oxidize and remove the free fatty acids released by the lysosome.

Cellular mitochondrial health relies on mitophagy, a form of selective autophagy whereby autophagosomes target damaged or dysfunctional mitochondria for degradation within lysosomes^12,13^. Damaged mitochondria not only have inefficient energy production but are also a major source of reactive oxygen species (ROS)^14^, which can ultimately lead to cell death^15^. Mitophagy is central to the maintenance of mitochondrial homeostasis, the promotion of optimal energy production, and the protection against oxidative stress^16^. Impaired mitophagy disrupts efficient fatty acid oxidation and elevates oxidative stress, and thus has been shown to promote MASLD onset and progression to MASH^17,18^.

Both mitophagy and lipophagy rely on functional lysosomes^19^. There has been extensive study on many aspects of lysosomal biology where disruption of characteristics such as the acidic environment, the activity of lysosomal hydrolases and lipases, or the regulation of lysosome biogenesis can lead to diseases such as lysosomal storage disorders^20^. A less studied aspect of the lysosome is its positioning. Lysosomes are typically distributed throughout the cell; however, they must travel to the perinuclear region to fuse with autophagosomes^21^. We and others have reported that transmembrane protein 55B (TMEM55B), also known as phosphatidylinositol-4,5-biphosphate 4-phosphatase 1 (PIP4P1), alters multiple aspects of lysosome biology including lysosome intracellular positioning and mobility^22–24^. TMEM55B induces dynein-dependent lysosomal retrograde trafficking by binding and recruiting JIP4 (C-Jun-amino-terminal kinase-interacting protein 4), a dynein motor, to the lysosomal surface^22,25^. In the context of fatty acid incubation, we previously reported that inhibition of TMEM55B in hepatoma cell lines led to perinuclear localization of lysosomes, which appeared both enlarged and immobile^24^. In addition, a recent report further demonstrated the importance of TMEM55B as a molecular sensor that links autophagic flux and lysosomal repair during oxidative stress^26^. To further this line of investigation, here we sought to test the impact of TMEM55B on lipophagy and mitophagy. Given the established role of lipophagy and mitophagy on MASLD, we also sought to determine whether TMEM55B impacts MASLD onset and progression.

## MATERIALS AND METHODS

### Quantitative real-time PCR

RNA was extracted from mouse primary hepatocytes, livers, skeletal muscles, or adipose tissues, and reverse transcribed into cDNA as previously described^27^. All assays were performed in triplicate using 100ng cDNA on an ABI PRISM 7900 Sequence Detection System using TaqMan and SYBR Green qPCR assays. *TMEM55B* was quantified using a Taqman primer (Hs00292741 for human and Mm01319582_m1 for mouse) from Thermo Fisher Scientific, and *Jip4* SYBR Green primers are as follows (Forward: GGCGGCTCGAGAAAATCCGTTCTA; Reverse: AATGCGGCCGCAACTCAATCAAC). Real-time PCR results were normalized to *CLPTM* (Hs00171300, Thermo Fisher Scientific, Waltham, MA) as an internal control.

### Western blot analyses

Tissues were homogenized in RIPA lysis buffer with Protease Inhibitor Cocktail (Thermo Fisher Scientific), centrifuged for 15 min at x16,000g at 4°C, and the supernatants were collected. Samples were eluted with Laemmli buffer and denatured at 95°C for 5 min before loading to 8% or 4-20% Tris-Glycine gels (Thermo Fisher Scientific). After transferring, PVDF membrane was incubated with anti-Tmem55b or anti-Gapdh antibody at 4°C overnight, washed 3 times and incubated with secondary antibody for 1 hr at room temperature. Enhanced chemiluminescence substrate (Thermo Fisher Scientific) was used for protein detection. Quantitative analysis of protein bands was performed using Image J (NIH, Bethesda, MD).

### Cell culture and transfections

HepG2 or Huh7 cells were grown at 37°C and 5% CO2 in Eagle’s Minimum Essential Medium (EMEM) (ATCC, Manassas, VA) supplemented with 10% FBS (HyClone, Logan, Utah), 500 U/mL penicillin/streptomycin, and 2 nmol /L GlutaMAX (Invitrogen). *TMEM55B* and/or *JIP4* knockdown was achieved by transfection of 80,000 HepG2 cells/well in 12-well plates using either siRNAs (Life Technologies) targeting *TMEM55B* (S40499), *JIP4* (S17233) or non-targeting control using Lipofectamine RNAimax (Invitrogen) with 20 nM siRNA for 48 hours as previously described^24^. For double knockdown, siRNAs targeting *TMEM55B* and *JIP4* were added at the same time. Cellular phenotypes were quantified 48 hours post-transfection.

### Creation of *JIP4* Knockout (KO) cells

To create *JIP4* KO cell lines, single gRNA targeting *JIP4* (VB211203-1192efm, Supplemental Fig.S7A) and non-targeting gRNA as negative control (VB220208-1241bcc, Supplemental Fig.S7B) were designed and purchase from VectorBuilder (Chicago, IL). Cas9 2NLS nuclease is purchased from Synthego (Redwood City, California). Cells were transfected with 5 μg negative control or Jip4-Cas9/sgRNA complex using Cell Line Nucleofector™ Kit T (VCA-1002, Lonza) with Amaxa Biosystems Nucleofector II. After transfection, cells were grown in media with 500 µg/ml G418 for 10 days, and 200 µg/ml G418 for another 4 days to select for plasmid expressing cells.

### Flow cytometry

General lysosomal activity was quantified using the Lysosomal Intracellular Activity Assay Kit (ab234622, Abcam). Briefly, cells were incubated with a lysosome-specific self-quenched substrate for 1 hr at 37°C and 5% CO2. The fluorescence signal (generated by the substrate degradation in the lysosome) is proportional to the intracellular lysosomal activity. Bafilomycin A1, which inhibits lysosomal V-ATPase and prevents lysosomal acidification^28^, was used as a control. Lysosomal acid lipase activity was quantified using the LysoLive™ Lysosomal Acid Lipase Assay Kit (ab253380, Abcam). Cells were incubated with LipaGreen^TM^, a substrate that becomes fluorescent upon cleavage by lysosomal acid lipase. To measure mitochondrial membrane potential, HepG2 cells were harvested with 0.05% Trypsin-EDTA and resuspended in PBS with 50 nM tetramethylrhodamine ethyl ester (TMRE, T669, Invitrogen) for 25 minutes at 37°C and 5% CO2. To measure exogenous fatty acid uptake, cells were incubated with 200 ng/ml BODIPY493/503 (D3922, Invitrogen), 1 µm BODIPY-labelled C12-(D3822, Invitrogen), 2 µm BODIPY™ 558/568 C12 (D3835, Invitrogen) or 1 µm C16-FA (D3821, Invitrogen) for 30 mins, washed 3 times with PBS, harvested with 0.05% Trypsin-EDTA. For all assays, fluorescence intensity was quantified by BD FACS Calibur flow cytometer as the median fluorescence values of 10,000 gated events.

### Lipase assay

Cellular lipase activity was measured using the Lipase Assay Kit (Colorimetric) (ab102524, Abcam). Briefly, after harvesting, cells were homogenized, and incubated with reagents for 60 – 90 minutes at 37°C and absorbance (OD570 nm) was measured in the kinetic mode using BioTek Synergy H1 Plate Reader (Agilent Technologies). Lipase activities were calculated as nmol/min/mL based on the standard curve.

### Immunofluorescence staining and confocal microscopy

For lysosome staining, cells were seeded on glass slide coverslips within 12-well plates at a density of 80,000, incubated with 75nM LysoTracker DND-99 probes for 1 hr at 37°C and 5% CO2, washed with PBS and fixed with 4% paraformaldehyde in PBS for 10 min at room temperature. Cells were then permeabilized with 0.25% Triton-100 in PBS for 10 min at room temperature, washed with PBS, and incubated with 1% BSA, 22.52 mg/mL glycine in PBST (PBS+ 0.1% Tween 20) for 30 min to block unspecific binding of the antibodies. Cells were incubated with primary antibodies (i.e. LAMP1, PLIN2, LC3B) diluted in 0.1% PBST at 4°C overnight. After 3 washes with PBS, cells were incubated with secondary antibodies diluted in 0.1% PBST for 1 h at room temperature, washed 3 times with PBS, and mounted with ProLong™ Gold Antifade Mountant with DNA Stain DAPI (P36935, Invitrogen). For intracellular lipid accumulation, cells were incubated with 1µM oleate for 24hr, stained with 100 μg/mL Nile red (72485, Sigma) for 30 min, and fixed with 4% paraformaldehyde. All samples were examined under a Zeiss LSM 710 confocal laser-scanning microscope equipped with X63 oil-immersion objective. For quantitative image analysis, 10–16 randomly chosen fields that included 1–5 cells each were scanned using the same setting parameters (i.e., laser power and detector amplification) below pixel saturation. The mean intensity per field was determined using the histogram function in the Zeiss LSM 710 Software, and pixel values above background levels were quantified. All the experiments were repeated at least three times, and representative images are shown. To quantify colocalization (i.e., PLN2/LysoTracker, C16/LysoTracker, C12/LysoTracker, C12/Lamp1, C12/MitoTracker, C12/LDs, C12/LC3B colocalization), 10–14 randomly chosen fields that included 1–5 cells each were scanned and analyzed in Carl Zeiss software AIM using the Pearson’s Correlation Coefficients. All imaging parameters and analyzing settings remained the same for all data acquisition within one experiment.

### Fluorescent FA Pulse-Chase

Cells were incubated with EMEM with 10% fetal bovine serum,2mM glutamax, and 500 U/mL penicillin/streptomycin) containing 1 mM BODIPY 558/568 C12 (Red C12, D3835, Invitrogen) or 2 mM BODIPY FL C12 (Green, D3822, Invitrogen) for in duplicate wells to allow the fluorescent lipids to incorporate into LDs. After 16 hr, one well was collected as timepoint “HR 0”. The other well was washed three times with PBS, chased with DMEM, and collected after 24 hr as timepoint “HR 24”. At the time of collection, cells were washed twice with PBS, fixed with 4% paraformaldehyde in PBS for 10 min, and permeabilized with 0.25% Triton-100 in PBS for 10 min at room temperature. Mitochondria were labeled with 100 nM MitoTracker Deep Red FM for 30 min, LDs were labeled with 1µM BODIPY 493/503 for 30 min, and lysosomes were labeled with 75 nM LysoTracker Red DND-99 for 1 hr. All cells were imaged with Zeiss LSM 710 confocal laser-scanning microscope with X63 oil-immersion objective, and analyzed as described in the confocal microscopy section above.

### Live cell imaging

Cells were incubated with 0.2 μM LysoTracker (DND-99, Life Technologies) for 15min, 1 μM BODIPY C12 (Green, Life Technologies) for 30 minutes, and 0.2 μM MitoTracker Deep Red (Life Technologies) for 15 minutes and imaged on a spinning disk confocal microscope (Nikon CSU-X1 dual camera platform equipped with Okolab stagetop incubation system and an iXon Ultra 888 EMCCD camera) using a 100X Apo TIRF. Imaging data was collected in 10–20 movies per condition with the same laser power, exposure, and electron-multiplying gain settings for all conditions. Images were subjected to Gaussian filtering and background subtraction in Imaris v 9.6 (Bitplane, Concord, MA). For analysis of trafficking parameters, labeled vesicles were subjected to surface reconstruction using the Surfaces and Tracks modules, and track length, track displacement, and track lifetimes were calculated using Imaris. For analysis of mitochondrial volume, MitoTracker-labeled mitochondria were subjected to surface reconstruction in Imaris, and automated segmentation by color-coding based on the volume of the connected components was used for 3D surface rendering of mitochondria.

### Transmission electron microscopy

Cells were grown on MatTek glass bottom dishes (P35G-1.5-14-C, MatTek), fixed in 2% glutaraldehyde and 2% paraformaldehyde solution for 24 hrs and washed 3 times for 5 min each in 0.1 M sodium cacodylate buffer, pH 7.4. Samples were then post-fixed in 1% osmium tetroxide with 1.6% potassium ferricyanide (KFECn) in 0.1 M sodium cacodylate buffer for 30 min, and washed 3 times for 15 min each with PBS. Cells were dehydrated in a serial diluted ethanol solution of 30, 50, 70, 90, and 100%, for 10 min each and infiltrated with 50% Epon-Araldite resin (containing benzyldimethylamine accelerator) for 1hr, followed by 100% resin for 1 hr. Excess resin was removed from the MatTek dishes containing cells and polymerized at 60°C for 48 hrs. Using a dissecting blade, cells embedded in resin were mounted on resin-embedded blocks and serial sections of 70-150 nm thickness were cut on a Reichert-Jung Ultracut E microtome and set on 1 x 2-mm slot grids covered with 0.6% Formvar film. Sections were then post-stained with 1% aqueous uranyl acetate for 7 min and lead citrate for 4 min. Samples were imaged on an FEI Tecnai 12 transmission electron microscope equipped with a 2k x 2k CCD camera with a 40 Megapixel/sec readout mode. Images were analyzed using ImageJ software according to the method by Lam et al^1^.

### Measures of mitochondrial levels and oxidative stress

HepG2 cells were transfected with siRNAs targeting *TMEM55B* or a scrambled control siRNA, and plated in 96-well plates at a density of 1×10^e4^ per well. After 48 hours, cells were incubated with 200 nmol MitoGreen to stain mitochondria or 5 μmol CellRox Green to measure cellular oxidative stress for 30 mins, or 5 μmol MitoSox Red in HBSS (14025076, Gibco) for 10 min to measure mitochondria-specific oxidative stress, and 5µg/ml Hoechst for 15 min to stain the nuclei. After 3 washes with PBS, fluorescence levels were quantified with BioTek Synergy H1 Plate Reader (Agilent Technologies). To visualize MitoSox staining, transfected cells were also plated in 6-well plates at a density of 2×10^e5^ per well. After 48 hours, cells were treated with 5 μmol MitoSox Red in HBSS (14025076, Gibico) for 10 min, and imaged with Keyence at 20X.

### Measures of mitochondria function

10,000 HepG2 cells or 7,000 mouse primary hepatocytes were seeded into each well of a Seahorse Bioscience (Agilent, Santa Clara, CA) tissue culture 96-well plate coated with or without poly-D-Lysine and incubated overnight at 37 °C in 5% CO_2_. Cells were washed twice with DMEM assay media (sodium bicarbonate- and glucose-free DMEM supplemented with glutamine and penicillin/streptomycin, pH 7.4) and incubated for 1 hr at 37 °C without CO_2_. Measures of mitochondrial function and oxidation of exogenous and endogenous fatty acids were determined by recording oxygen consumption rates (OCR, pmol/min) on a Seahorse Bioscience XFe96 extracellular flux Analyzer. After measuring basal respiration, the injection of oligomycin was used to measure ATP production (final concentration 2 μM), and the injection of fluorocarbonyl cyanide phenylhydrazone (FCCP) was used to detect maximal respiration (final concentration 2μM). The final injection of antimycin A and Rotenone was to measure spare respiratory capacity and non-mitochondrial respiration. Background values were determined from cell-free wells and subtracted from sample values. Cellular protein levels were determined using a Bradford protein assay. Data were normalized to total protein in each well. To measure fatty acid β-oxidation rates, cells were seeded in XF96 well plates and starved overnight in substrate-limited DMEM supplemented with 0.5 mM of glucose, 1 mM of glutamine, 0.5 mM of carnitine, and 1% FBS. The bovine serum albumin (BSA)-conjugated palmitate (55.8 nM, Seahorse Bioscience) was added to a final concentration of 10 nM, and basal OCR of cells treated with palmitate-BSA or BSA vehicle alone were measured. For cell permeabilization, 1 nmol of Seahorse XF Plasma Membrane Permeabilizer (102504-100, Agilent Technologies) was used.

Mouse liver mitochondria were isolated and quantified as described^29^. Briefly, liver tissues were harvested and minced in cold MSHE buffer (210 mM Mannitol, 70 mM sucrose, 5 mM HEPES, 1 mM EGTA, pH 7.2) + 0.5%BSA, and homogenized with gentleMACS™ Dissociator (Miltenyi Biotec). The homogenate was centrifuged at 4°C at 800g for 10 min, after which the supernatant was transferred to another tube, and centrifuged again at 4°C at 8000g for 10 min. The pellet was resuspended in 800µl of MSHE buffer + 0.5%BSA and centrifuged again at 4°C at 8000g for 10 min. The pellet was resuspended in 30µl of MAS buffer (220 mM mannitol, 70 mM sucrose, 10 mM KH2PO4, 5 mM MgCl2, 2 mM HEPES, 1.0 mM EGTA, 0.2 % BSA, pH 7.2). Protein concentration was quantified with Pierce™ Bradford Plus Protein Assay Kits (23236, Thermo Scientific). 4 µg of mitochondria were plated each well in a XF96 well plate. OCR was measured after addition of 4 mM ADP (Complex V substrate), oligomycin, FCCP and antimycin as describe above. OCR was normalized per μg mitochondrial protein.

### Animal studies

Animal studies were performed with approval and in accordance with the guidelines of the Institutional Animal Care and Use Committee at the University of California San Francisco (UCSF). Animals were cared for according to the recommendations of the Panel on Euthanasia of the American Veterinary Medical Association. The animal facility is Association for Assessment and Accreditation of Laboratory Animal Care (AAALAC) approved and is responsible for the health and husbandry of animals. Animal studies comply with the Animal Research: Reporting of In Vivo Experiments (ARRIVE) guidelines. Mice were housed in a climate-controlled Department of Laboratory Animal Medicine facility with a 12-hour light-dark cycle and ad libitum access to food and water.

### *Tmem55b* knockout mouse model

*Tmem55b* floxed mice were purchased from Cyagen Biosciences who inserted *loxP* sites flanking *Tmem55b* exons 1 to 6 in a C57BL6/N background. The *Tmem55b^fl/+^* mice were backcrossed 7 times to the C57BL/6J strain, and subsequently crossed with Sox-2-Cre mice (Strain #:008454, JAX) on a C57BL/6J background to generate *Tmem55b^+/−^* mice. These mice intercrossed to generate whole-body *Tmem55b* knockout and littermate control *Tmem55b^+/+^* mice. Mice were confirmed to have a >99% C57BL/6J background using The Jackson Laboratory’s C57BL/6 Substrain Characterization Panel. Primary hepatocytes were isolated from 10-week old male and female mice at the University of California, San Francisco Liver Center using the standard collagenase method^30^.

6-week old mice were fed a GAN diet (40% kcal fat, 20% kcal fructose, 2% cholesterol) for 21 weeks, after which mice were euthanized after 4 hr fast. Blood was collected via cardiac puncture, and plasma isolated by centrifugation at 850× g for 15 min at 4 °C. Tissues were flash frozen in liquid nitrogen, and a portion of liver was fixed in 10% formalin, washed with PBS the next day, and stored in 70% ethanol for histology.

### ASO-mediated *Tmem55b* knockdown mouse model

C57BL/6J male and female mice were purchased from Jackson Laboratory (Bar Harbor, ME). 6-week-old animals were i.p. injected with 25 mg/kg body weight/week of antisense oligonucleotides (ASO) targeting *Tmem55b* or a non-targeting control (Ionis Pharmaceuticals) and fed a GAN Diet. Body weight was measured prior to the first ASO injection, and weekly until sacrifice. After 7, 21, or 29 weeks, mice were fasted for 4 hours, euthanized and blood and tissues collected as described for the *Tmem55b* knockout model.

### Hepatic lipid measurements

Liver tissues were homogenized with GentleMacs (Miltenyi Biotec Inc. Auburn, CA) and lipids extracted with chloroform-methanol (2:1) according to the Folch method^31^. Chloroform extracts were dried under N_2_ gas and resuspended in 200 μl isopropyl alcohol containing 10% Triton X-100. TAG was measured using the L-Type TG M kit (Wako Chemicals, Richmond, VA) following manufacturer’s instructions.

### Mouse liver Immunohistology

For Oil red O (ORO) staining, mouse liver was embedded in Tissue-Tek, sectioned, and stained in 0.5% Oil Red O solution in propylene glycol for 30 min. The slides were transferred to a 85% propylene glycol solution for 1 min, rinsed in distilled water 2 times and processed for hematoxylin counter staining. For H&E, Sirius Red, and Masson Trichrome staining, a portion of mouse liver was fixed with 10% formalin, embedded in paraffin, and cut into 4-µm sections. Sections were deparaffinized with xylene, rehydrated using an ethanol gradient (100, 100, 95, 95, and 80%), and stained with different dyes. All slides were assessed by a pathologist blinded to the sample identity.

### Plasma aminotransferase measurements

Mouse plasma aspartate aminotransferase (AST) and alanine aminotransferase (ALT) levels were measured by enzymatic end point measurements using enzyme reagent kits (Ciba-Corning Diagnostics Corporation) in an AMS Liasys 330 Clinical Chemistry Analyzer.

### RNA-sequencing

Total RNA was extracted from livers of male mice treated with *Tmem55b* (N=4) or non-targeting control (N=4) ASOs and fed a Western diet for 4 weeks, checked for quality on a Bioanalyzer, and made into polyA-selected, strand-specific RNA-seq libraries for 150 bp paired-end sequencing on Illumina NovaSeq machines. Sequence fragments were aligned to the mouse GRCm39 genome and GENCODE transcriptome using STAR 2-pass alignment^32^. Fragments aligning to annotated genes were counted using featureCounts^33^ and adjusted for library size using DESeq2^34^. DESeq2 was used for differential expression analysis, and p-values were adjusted for multiple testing using a false discovery rate (FDR) approach.

### Statistics

For *ex vivo* and *in vitro* experiments, representative results are reported. Data are shown as mean ± standard error of the mean (SEM). Grubb’s test for outliers was used to identify statistical outliers. Continuous variables for two groups were compared using Student’s t-tests. Continuous variables for more than two groups were compared using one-way analysis of variance (ANOVA) with Tukey’s post hoc test. Analyses were performed using GraphPad Prism 7 software (GraphPad Software, Inc. La Jolla, CA, USA). P values <0.05 were considered statistically significant.

## RESULTS

### Loss of *TMEM55B* accelerates diet-induced MASLD onset and progression in mice

To assess the effect of TMEM55B inhibition *in vivo,* we injected Western diet-fed C57BL/6J male mice with ASOs targeting *Tmem55b* or non-target control (NTC) (**Supplemental Fig. S1A**). After 6 weeks, *Tmem55b*-ASO treated male animals had significantly greater hepatic lipid accumulation as observed by Oil Red O and H&E staining (1.5-fold, **Fig. 1A, 1B**), and through direct quantification of extracted hepatic lipids (1.4-fold, p<0.0001, **Fig. 1C**). Next, we tested whether *Tmem55b* knockdown accelerated MASLD progression in C57BL/6J male mice fed the metabolic dysfunction associated steatohepatitis (MASH)-inducing GAN diet (40% kcal fat, 20% kcal fructose, 2% cholesterol) (**Supplemental Fig. S1B)**. After 21 weeks, *Tmem55b*-ASO treated animals had a higher level of fibrosis compared to NTC treated animals (p<0.05, **Fig. 1D**), a difference that was even more pronounced after 29 weeks (p<0.01). These findings were consistent with our RNAseq analysis of 4-week Western diet and *Tmem55b* vs. NTC ASO treatment (GEO# GSE273884) which demonstrated a functional enrichment of upregulated genes involved in cell adhesion (*Flrt2*, *Egfl7*, *Col8a1*, *Pkp3*, *Pcdh17, Mcam*) in the liver (**Supplemental Fig. S1C**).

**Figure 1.**
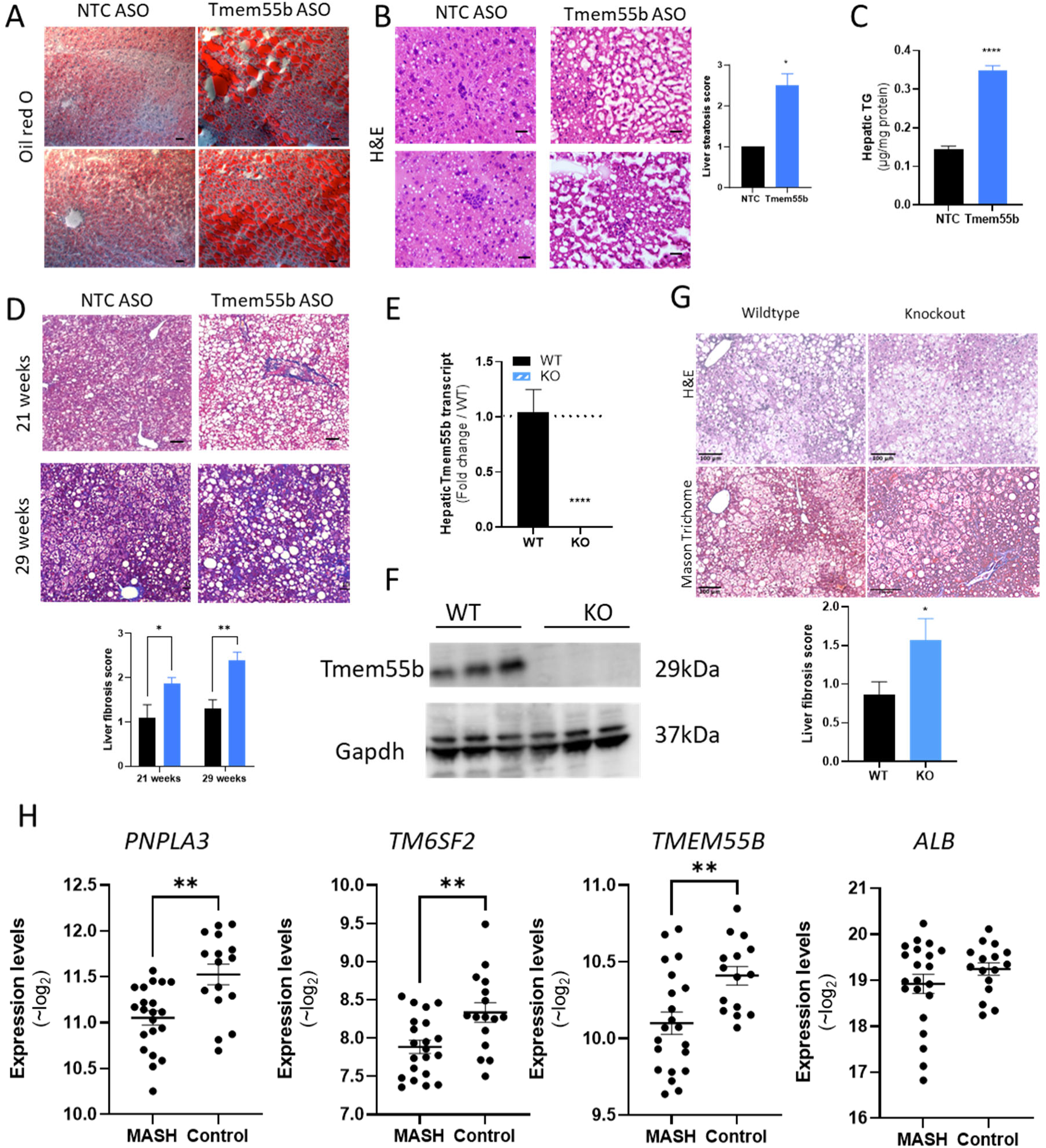
Loss of Tmem55b led to hepatic lipid accumulation and NASH development. Six-week old male C57BL/6J mice were treated with an ASO against Tmem55b (Tmem55b-ASO) or a non-targeting control (NTC) at a dose of 25 mg/kg body weight/week and fed a Western diet (0.2% cholesterol, 42% fat) for 6 weeks. Liver tissues were stained with Oil Red O (A), H&E (B) (Scale bars=10 μm), or homogenized and lipid was extracted, and triglyceride levels were measured (C). D) Male C57BL/6J mice were treated with Tmem55b-ASO or NTC-ASO and fed Western diet for 21 or 29weeks. Mouse liver was stained with Masson Trichrome. Liver fibrosis, the NAFLD activity score (NAS), % macrovesicular steatosis and inflammation were assessed by a blinded pathologist (N=5-12). E) Hepatic *Tmem55b* transcript and (F) protein levels were quantified by qPCR and western blot in wildtype and TMEM55B knockout mice. G) Six-week-old male wildtype and TMEM55B knockout mice were put on GAN diet for 12 weeks. Mouse liver was stained with H&E and Masson Trichrome. Liver fibrosis was assessed by a blinded pathologist (Scale bars=100 μm). N=4-10. For the animal studies, quantified results are presented as mean ± s.e.m. *p<0.05, **p<0.01, ***p<0.001, ****p<0.0001 vs. NTC or Tmem55b+/+ by one-way ANOVA or Student’s t-test. H) TM6SF2, TMEM55B and ALB transcript levels in iPSC-Heps from NASH cases (n=21) and controls (n=15) quantified by RNAseq. Variance stabilized (∼log2) values were calculated in DESeq and plotted. **p<0.005.

As there may be non-specific effects of ASOs, we also created a *Tmem55b* knockout model (as described in **Supplemental Fig. S1D**), where we confirmed the absence of hepatic *Tmem55b* transcript (**Fig. 1E**) and protein (**Fig. 1F**). Male and female mice were fed a GAN diet starting at 6-weeks-old for 12-weeks (**Supplemental Fig. S1E**). Compared to wildtype littermates, both male and female *Tmem55b* KO animals had significantly increased liver fibrosis (p<0.05) (**Fig. 1G, Supplemental Fig. S2A**). We also observed increased liver weight (p<0.01) and trends of greater plasma ALT and hepatocyte ballooning in the male *Tmem55b* KO (**Supplemental Fig. S2B**).

### *TMEM55B* levels are lower in iPSC-derived hepatocyte-like cells from MASH patients compared to healthy controls

Transcript differences detected in livers from MASLD cases and controls may be consequence of, rather than a cause of, the disease. This is particularly salient for *TMEM55B*, which is regulated by cholesterol levels^27^. In contrast, differences observed in induced pluripotent stem cell derived hepatocyte-like cells (iPSC-Heps) may reflect genetic (and thus potentially causative) contributors to disease. RNAseq data was available (GEO# GSE138312) from a cohort of iPSC-Heps from biopsy-defined MASH cases and healthy controls. Cohort details have been previously reported^35^. Both *PNPLA3* and *TM6SF2* are known determinants of MASLD, where loss of function alleles in humans and hepatic knockout in murine models promote MASH development^36,37^. As expected iPSC-Heps from MASH cases had lower transcript levels of *PNPLA3* and *TM6SF2* compared to controls (**Fig. 1H**). In addition, there were similar expression level of albumin (*ALB*), a hepatocyte-specific gene, indicative of equivalent differentiation efficiency between the two groups. Importantly, we found reduced *TMEM55B* transcript levels in the MASH iPSC-Heps (P<0.01, **Fig. 1H**), while correlation does not equal causation, these findings are consistent with the likelihood that reduced levels of TMEM55B may lead to hepatic steatosis and MASH.

### *TMEM55B* knockdown increases autophagy, specifically lipophagy

TMEM55B is a transmembrane protein located on late endosomes and lysosomes^38^. We previously reported that *TMEM55B* knockdown in HepG2 cells increased levels of the lysosome marker lysosome-associated protein 1 (LAMP1) and led to lysosomal clustering in the perinuclear region upon exposure to fatty acids^24^. Here we have extended these observations in primary murine hepatocytes from *Tmem55b* knockout (KO) and wildtype (WT) male mice (**Supplemental Fig. S1D**). After incubation with 1mM oleate for 1 hour, lysosomal volume was 71% higher in hepatocytes from the KO vs. WT mice (p<0.05) while there was no difference in lysosome number (**Fig. 2A)** with live cell imaging. Similarly, increased lysosomal areas without changes in lysosome number were observed with confocal microscope in HepG2 cells treated with a siRNA targeting *TMEM55B* compared to scrambled (scr) siRNA controls (**Supplemental Fig. S3A, S3B**), and decreased lysosome track speed, length and duration was also observed with live cell imaging (**Supplemental Fig. S3C**), indicating decreased lysosome mobility with *TMEM55B* knockdown. Consistent with the greater lysosome volume, *TMEM55B* knockdown in HepG2 cells led to increased lysosomal intracellular activity by 24% (p<0.05, **Fig. 2B**), and a 33% (p<0.001) increase of lysosome acid lipase, which is responsible for hydrolyzing lysosomal triglycerides and cholesteryl esters^39^ (**Fig. 2C**). Bafilomycin A1 serves as a positive control here, which inhibits lysosomal V-ATPase and prevents lysosomal acidification^28^, resulting in a decreased lysosomal activity (p<0.05, **Fig. 2B**).

**Figure 2.**
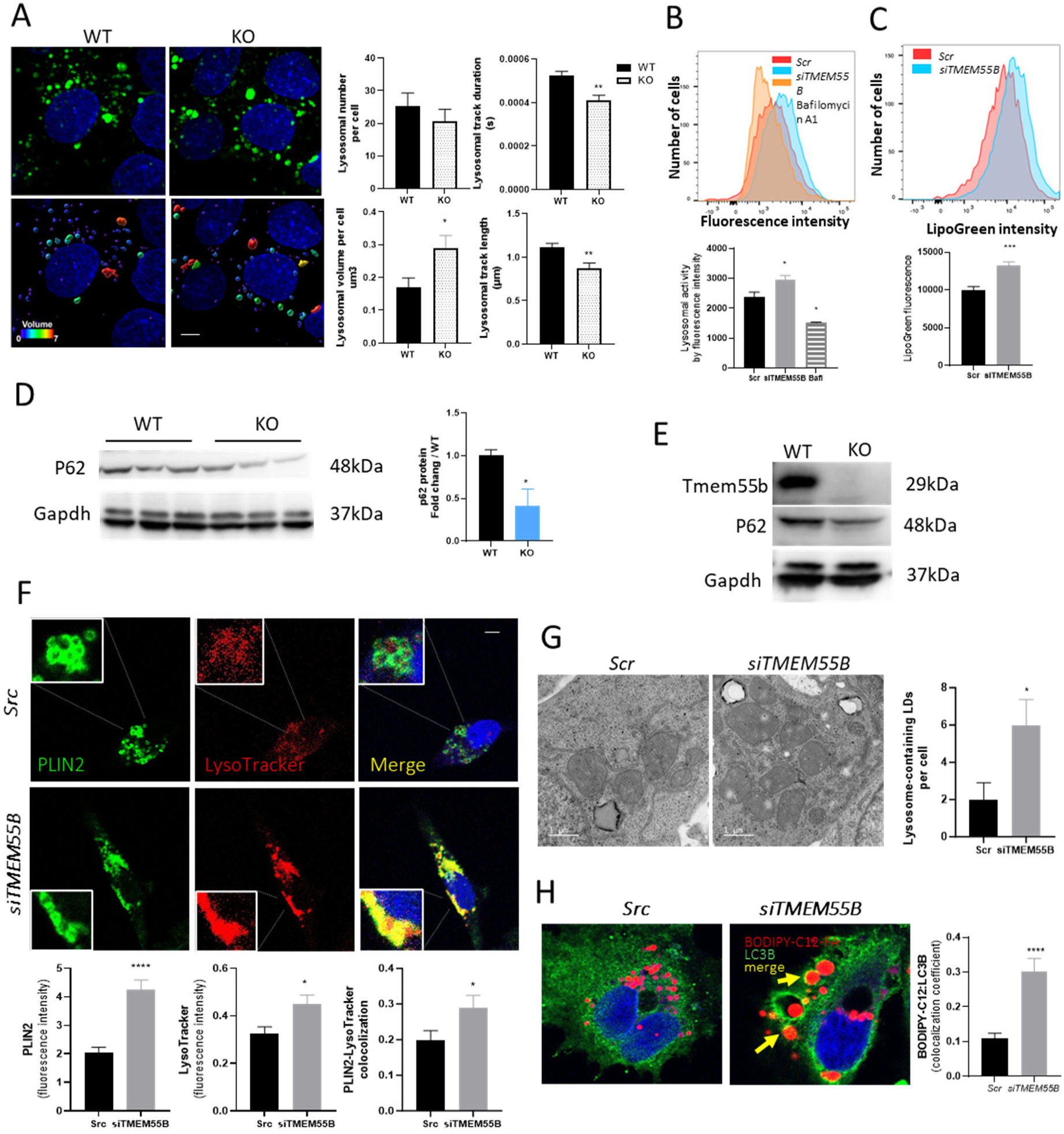
Tmem55b deficiency increased autophagy flux and lipophagy. A) Primary hepatocytes were isolated from Tmem55b knockout (KO) and Tmem55b wildtype (WT) male mice and treated with 1mM oleate for 1 hour before being stained with LysoTracker and imaged with Nikon spinning disk confocal microscope at100X (Scale bars=5 μm). For analysis of lysosome volume, lysosomes were subjected to surface reconstruction in Imaris, and automated segmentation by color-coding based on the volume of the connected components was used for 3D surface rendering of lysosomes (bottom panels). Representative images are shown here. HepG2 cells were transfected with TMEM55B or Scr siRNAs. B) Lysosome general activity and C) Lysosomal Acid Lipase activity were measured through flow cytometry. Six-week-old male wildtype and TMEM55B knockout mice were put on GAN diet for 12 weeks. and p62 protein levels were detected by immunoblot in mouse liver tissues (D) or primary hepatocytes isolated from WT and Tmem55b knockout mice. F) Cells were labeled with anti-PLIN2 and LysoTracker and examined by confocal microscopy (63X, scale bars=10 μm). G) Electron micrographs of HepG2 cells showing increased numbers of lysosome-containing lipid droplets (LDs) with Tmem55b knockdown (63X, scale bars=10 μm). H) After transfection, HepG2 cells were given 1 μm of red BODIPY™ 558/568 C12-FA overnight, stained with anti-LC3B antibody for 1 hr at room temperature followed by 2nd Goat anti-rabbit IgG H&L (Alexa Fluor® 488) and examined by confocal microscopy. Representative images showing lipid droplets trapped in LC3B-labelled autophagosomes. Representative images are shown here. Results are presented as mean ± s.e.m. *p<0.05, **p<0.01, ***p<0.001, ****p<0.0001 vs Scr or WT by Student’s t-test.

Autophagy plays a pivotal role in lipid metabolism in mammalian hepatocytes^10^, and it relies on lysosomal activity to break down TG and release FAs^19^. p62 protein levels are a measure of autophagic flux^40^. Defective autophagy leads to p62 accumulation, while increased autophagy results in reduced p62 levels^41^. In the liver tissues (**Fig. 2D**) and primary hepatocytes (**Fig. 2E**) from male *Tmem55b* knockout mice we observed reduced p62 protein levels compared to WT littermate controls, indicative of enhanced autophagy flux. We also confirmed that there was no *Tmem55b* transcript or protein expressed in primary hepatocytes isolated from the *Tmem55b* KO animals (**Fig. 2E**, **Supplemental Fig. S3D)**.

Lysosome-mediated lipid turnover, known as lipophagy, is a process by which lipids stored in LDs are directly transported to the lysosome, or through an autophagosome-dependent event, for the liberation of fatty acids^42^. We examined the co-localization of lysosomes and lipid droplets in *TMEM55B* siRNA and Scr siRNA treated HepG2 cells. In the control treated cells, staining for perilipin-2 (PLIN2), a lipid droplet membrane marker^43^, revealed distinct circles around the periphery of lipid droplets as expected (**Fig. 2F**). Upon *TMEM55B* knockdown, PLIN2 levels were increased, the peripheral circular structure was lost, and there was greater co-localization with lysosomes (1.5-fold, p<0.05, **Fig. 2F**), indicative of the formation of lipid-filled lysosomes (aka lipolysosomes)^44^. We confirmed this increase in lysosomal lipid accumulation (**Supplemental Fig. S3E, S3F**) with trace amounts (1μM) of BODIPY labeled C16-FA or C12-FA, and stained with LysoTracker or LAMP1. Under all conditions, *TMEM55B* knockdown increased lysosome intensity levels (1.2-fold, p<0.01) and colocalization of neutral lipids and lysosomes (1.5-fold, p<0.001, **Fig. 2F, Supplemental Fig. S3E, S3F**). In addition, we observed changes in lipolysosome intracellular positioning. Under control conditions, colocalization between lipids and lysosomes appeared primarily at the cell periphery (**Supplemental Fig. S3F**, yellow arrows), versus within the perinuclear region in the *siTMEM55B* treated cells (green arrows). We confirmed the increase in lipolysosomes using electron microscopy, where the number of lysosomes containing lipid droplets was increased by 3-fold (p<0.05, **Fig. 2G**). Lastly, knockdown of *TMEM55B* led to greater co-localization C12-FA to microtubule-associated proteins 1A/1B light chain 3B (LC3B), (**Fig. 2H**), a marker of autophagy. Together, these findings suggest that the loss of *TMEM55B* increases lipophagy.

### *TMEM55B* knockdown increases lysosomal fatty acid release to mitochondria

Lysosomes play a critical role in lipid metabolism and MASLD onset^45–47^. Enhanced lipophagy leads to increased free fatty acid release from lysosomes, primarily to mitochondria for β-oxidation^10^, as well as to lipid droplets^48^ or the outside of the cells^49^. In addition to lipophagy, intracellular fatty acids can be hydrolyzed from triglycerides stored within lipid droplets through cytosolic lipolysis via lipases^50^. Therefore, we first examined cytosolic lipase activity upon *TMEM55B* knockdown and found no effect in either HepG2 or Huh7 cells (**Supplemental Fig. S4A, S4B**). Given the impact of TMEM55B on lipophagy, we sought to test whether loss of *TMEM55B* impacts fatty acid mobilization and trafficking between mitochondria, lipid droplets, and lysosomes, using a pulse-chase assay in HepG2 cells (**Fig. 3A, Supplemental Fig. S4C**). As oleic acid (C18) is one of the most abundant fatty acids in the diet and circulation^48,51^, we used BODIPY C12-FA as its overall length with the attached fluorophore is equivalent to C18-FA^52,53^. HepG2 cells were incubated with 2µM BODIPY C12-FA for 16 hours to allow incorporation into neutral lipids and storage within lipid droplets and chased with substrate-limited DMEM for 24 hours after which cells were labeled with markers for lipid droplets, lysosomes or mitochondria (**Fig. 3A**) and visualized with confocal microscopy.

**Figure 3.**
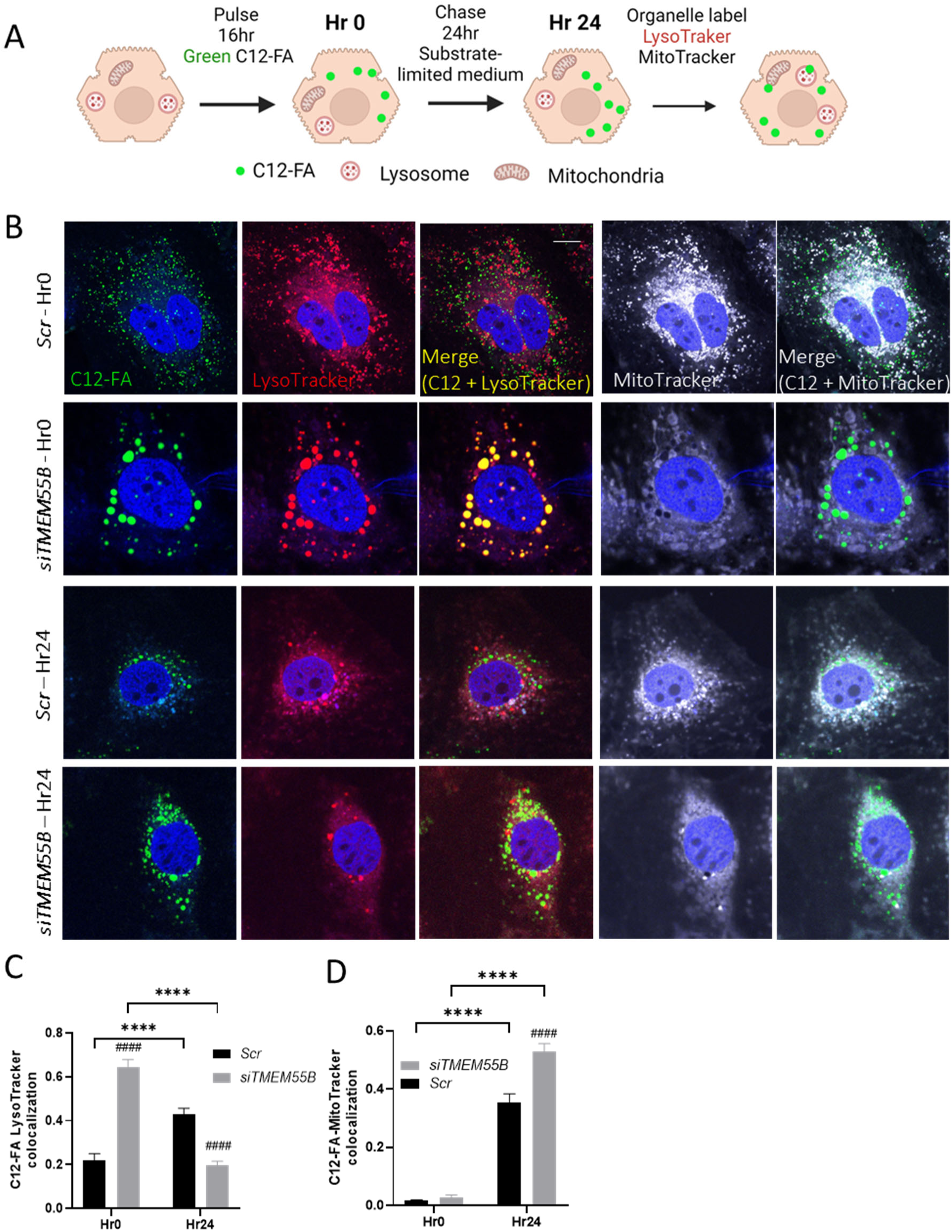
Elevated autophagy flux by TMEM55B knockdown increased lysosomal fatty acid release to mitochondria. A) Schematic representation of the FA pulse-chase assays. HepG2 cells were transfected with TMEM55B and Scr siRNAs, pulsed with Green C12-FA (2μM) overnight, washed, and chased with substrate-limited DMEM supplemented with 0.5 mM of glucose, 1 mM of glutamine, 0.5 mM of carnitine, and 1% FBS for 24 hours. After which, cells were labeled with LysoTracker Red and MitoTracker Deep Red, and imaged with Zeiss 710 confocal microscope at 63X (Scale bars=10 μm) (B). C12-FA localization to Lysosome and mitochondria (C) before and after the chase were quantified. (D). After incubation with 2μM BODIPY C12-FA for 30 mins, HepG2 cells were labeled with MitoTracker Red, imaged with Nikon spinning disk confocal microscope at 100X (Scale bars=3 μm), and FA-Mitochondria colocolization were quantified (mitochondria is red and FA is blue). Representative images are shown here. Results are presented as mean ± s.e.m. **p<0.01, ****p<0.0001 vs Hr0; ###p<0.001, ####p<0.0001 vs Scr by two-way ANOVA.

After the 16hr pulse (Hr0 of chase), *TMEM55B* knockdown led to greater localization of C12-FA to lysosomes (2-fold, p<0.0001, **Fig. 3B, 3C**) and lipid droplets (1.4-fold, p<0.0001, **Supplemental Fig. S4D**), suggestive of increased lipophagy and consistent with our prior observations of enlarged lipid-filled lysosomes (**Fig. 2F, Supplemental Fig. S3F**). In the control cells, after the 24-hour chase of substrate-limited media, we observed the expected increase of C12-FA colocalization to lysosomes and mitochondria, consistent with the known cellular response of increased fatty acid β-oxidation during starvation^54^. In addition, there was increased C12-FA colocalization to lipid droplets (**Supplemental Fig. S4D**), a widely reported paradoxical characteristic of hepatocytes in response to starvation^55^. However, in the *siTMEM55B* treated cells, after the 24hr chase, there was significantly greater C12-FA colocalization to mitochondria (1.5-fold, p<0.0001, **Fig. 3B, 3D**), and reduced colocalization to lysosomes compared to Scr controls (1.5-fold, P<0.0001, **Fig. 3B, 3C**). These findings show that loss of *TMEM55B* promotes both endogenous and exogenous fatty acid delivery to mitochondria.

### *TMEM55B* knockdown leads to mitochondrial dysfunction

Excessive fatty acid trafficking to mitochondria can lead to mitochondrial damage^56^, thus we next sought to test whether TMEM55B modulates mitochondrial function. Primary hepatocytes isolated from female *Tmem55b* KO animals showed dramatically reduced mitochondrial oxygen consumption rate (OCR) (**Fig. 4A**) compared to the ones from WT littermate controls. Basal respiration, ATP production, maximal respiration, non-mitochondrial respiration, proton leak, and spare capacity were all significantly decreased in *TMEM55B* knockout hepatocytes compared to WT controls (**Fig. 4B**). Similar reductions in mitochondrial activity were observed in HepG2 cells with TMEM55B knockdown (**Fig. 4C, 4D**). Notably, the effect of *TMEM55B* knockdown on OCR achieves levels essentially as low as the samples treated with Etomoxir (**Fig. 4C)**, a negative control which inhibits CPT-1, a FA transporter on the mitochondrial outer membrane^57^. While the addition of exogenous fatty acids (such as palmitate) normally increases both basal and maximal OCR, as seen in the control-treated cells, there was no change in OCR upon palmitate addition after *TMEM55B* knockdown, suggesting an inability to utilize exogenous fatty acid for mitochondrial β-oxidation (**Fig. 4A**). To exclude the possibility that the decrease OCR we observed is due to decreased FA uptake into the cells, we permeabilized HepG2 cells with XF Plasma Membrane Permeabilizer (PMP) and incubated with or without the addition of 40 µM palmitate-coA. Similar to the intact cells, the addition of palmitoyl-CoA increased OCR only in control but not *siTMEM55B* treated cells (**Supplemental Fig. S5A**). Lastly, we confirmed that *TMEM55B* knockdown prevents utilization of exogenous fatty acid in mitochondria isolated from male mice treated with an antisense oligonucleotides (ASO) against *Tmem55b* vs. non-targeting control (**Supplemental Fig. S5B**), a model that we previously showed to reduce hepatic *TMEM55B* transcript and protein levels by ∼60%^24^.

**Figure 4.**
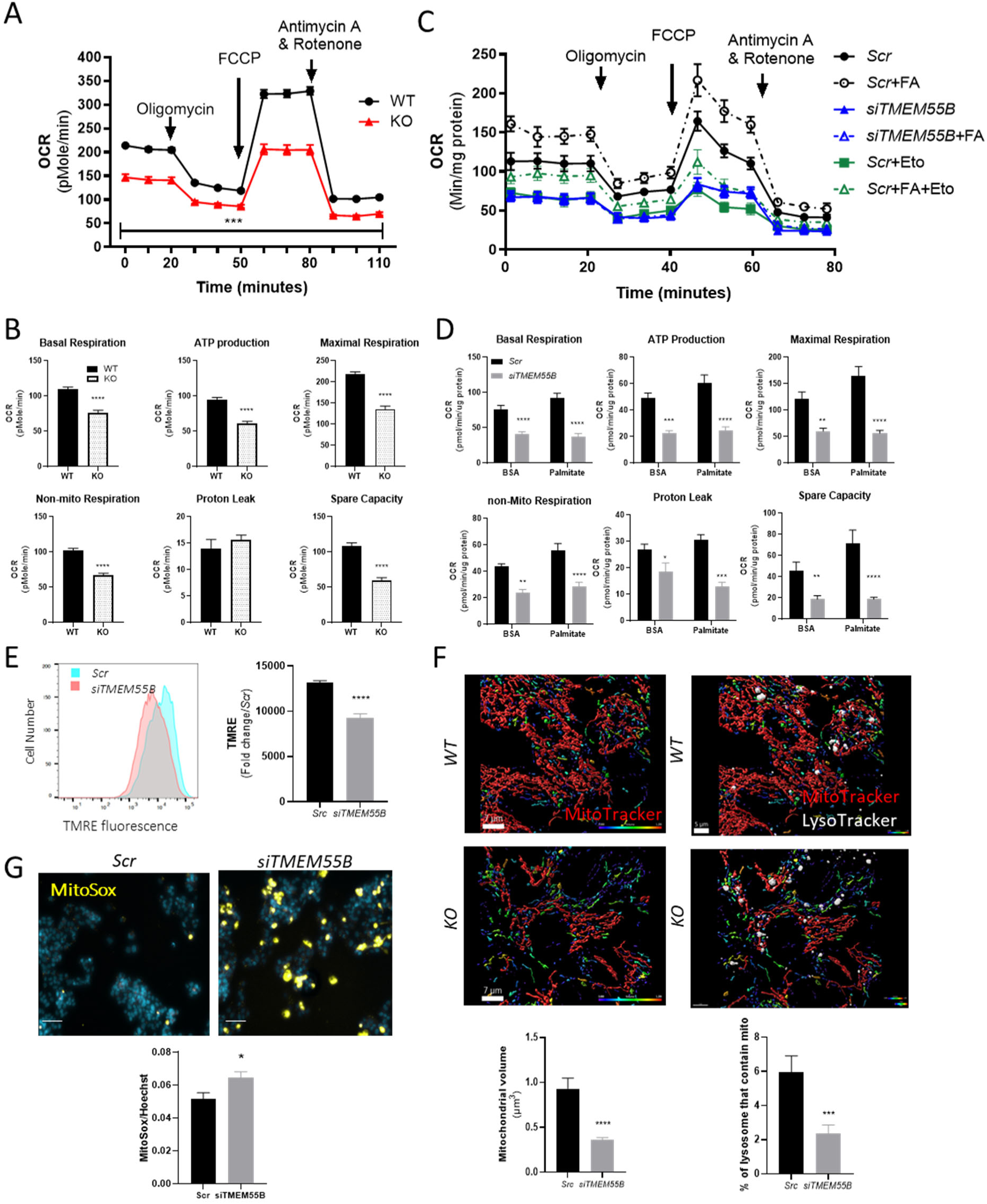
Tmem55b knockdown resulted in mitochondria dysfunction, increased oxidative stress and decreased mitophagy. A, B) OCR of primary hepatocytes from female wild type (WT) and Tmem55b−/− (KO) mice. C, D) HepG2 cells were transfected with siRNAs targeting TMEM55B (siTMEM55B) or scrambled (Scr) siRNA and oxygen consumption rate (OCR) was measured by Seahorse XF Analyzer with and without palmitic acid (125 mM, FA) and/or 40mM etomoxir (Eto). E) After transfection, mitochondrial membrane potential of HepG2 cells was measured by FACS quantification of TMRE staining. F) Primary hepatocytes were isolated from male WT and Tmem55b knockout mice, and the levels of mitochondrial dyes (MitoTracker) were quantified by live cell imaging with Nikon spinning disk confocal microscope at 100X (Scale bars=7 μm), and mitochondria (red) and lysosome (white) colocolization were visualized through live cell imaging and quantified (Scale bars=7 μm). In these images, red indicates healthy mitochondria, while unhealthy mitochondria are observed as blue or purple. G) mitochondrial oxidative stress was measured by MitoSox (Scale bars, 100 μm). For live cell imaging, Images were subjected to Gaussian filtering and background subtraction in Imaris v9.6. For analysis of mitochondrial volume, mitochondria were subjected to surface reconstruction in Imaris, and automated segmentation by color-coding based on the volume of the connected components was used for 3D surface rendering of mitochondria. Representative images are shown here. Results are presented as mean ± s.e.m. N=10-30 per treatment group. *p<0.05, **p<0.01, ***p<0.001, ****p<0.0001 vs Scr by Student’s t-test.

To determine whether the reduction in mitochondrial fatty acid β-oxidation could be due to impaired mitochondrial function or/and reduced levels of mitochondria, we next evaluated the effects of *Tmem55b* knockdown on mitochondrial membrane potential, essential for the production of ATP^58^. *TMEM55B* siRNA and control treated HepG2 cells were stained with TMRE (tetramethylrhodamine, ethyl ester), a dye that accumulates only in the membrane of polarized (active) mitochondria, and quantified with FACS. *TMEM55B* knockdown reduced TMRE levels by 35% (p<0.05, **Fig. 4E**). We also observed reduced mitochondrial volume by 61% (p<0.0001, **Fig. 4F**) with 3D-live cell imaging using MitoTracker Deep Red in primary hepatocytes from male Tmem55b knockout mice and WT controls, and decreased mitochondria intensity by 42% (p<0.001, **Supplemental Fig. S5C**) in fixed cells stained with MitoGreen in HepG2 cells.

Based on these findings, we further examined mitochondria morphology through live cell imaging and 3D reconstruction where we observed highly fragmented mitochondrial networks in primary hepatocytes from *Tmem55b* knockout mice compared to WT controls (**Fig. 4F**). Fragmented mitochondria are a sign of mitochondria dysfunction^59^. Typically, damaged mitochondria are degraded through mitophagy, a lysosome-mediated process^16^. One of the well-known regulators of mitophagy is p62^60,61^, which was decreased with *Tmem55b* knockout (**Fig. 2D, 2E**). Consistent with the reduced p62 levels, *TMEM55B* knockout led to a significantly lower percentage of lysosomes containing mitochondria (p<0.001, **Fig. 4F**), suggestive of impaired mitophagy. Mitochondrial damage often leads to oxidative stress from inefficient ATP production and increased production of reactive oxygen species^62^. Not surprisingly, *TMEM55B* knockdown led to significantly higher levels of oxidative stress using a generalized measure (CellRox, **Supplemental Fig. S5D**), and a specific measure of mitochondrial oxidative stress (MitoSOX™, **Fig. 4G, Supplemental Fig. S5E**) in HepG2 cells. Together, these findings suggest that loss of *TMEM55B* resulted in mitochondrial dysfunction possibly through overwhelming FA flux to mitochondria and/or impaired mitophagy.

### *TMEM55B* knockdown causes cellular steatosis through lysosome position

Given that the loss of *TMEM55B* led to an inability of cells to utilize exogenous FA for mitochondrial β-oxidation, we sought to test whether there were defects in FA uptake. We incubated primary hepatocytes (**Fig. 5A)** and HepG2 cells (**Fig. 5B)** with BODIPY-C12 for 30 minutes. To our surprise, there was greater FA uptake in primary murine hepatocytes from *Tmem55b* KO vs. WT mice (**Fig. 5A**) and HepG2 cells treated with *TMEM55B* vs. Scr siRNAs (**Fig. 5B**) using flow cytometry. *TMEM55B* knockdown led to greater FA volume/cell (p<0.05, **Fig. 5C**), indicative of increased FA uptake. *TMEM55B* knockdown also caused FA to move more slowly and travel over shorter track lengths (p<0.05, **Fig. 5C**). To test whether loss of *TMEM55B* also impacts trafficking of exogenous fatty acids to the mitochondria, we incubated HepG2 with BODIPY-C12 for 30 minutes and stained with MitoTracker Deep Red and found a 1.5-fold increase in FA-mitochondria colocalization upon *TMEM55B* knockdown (p<0.01, **Fig. 3D**). We found no differences in protein levels of CD36, an abundant hepatic FA transporter, in the livers between ASO-*Tmem55b* and ASO-NTC treated male or female mice (**Supplemental Fig. S6A**).

**Figure 5.**
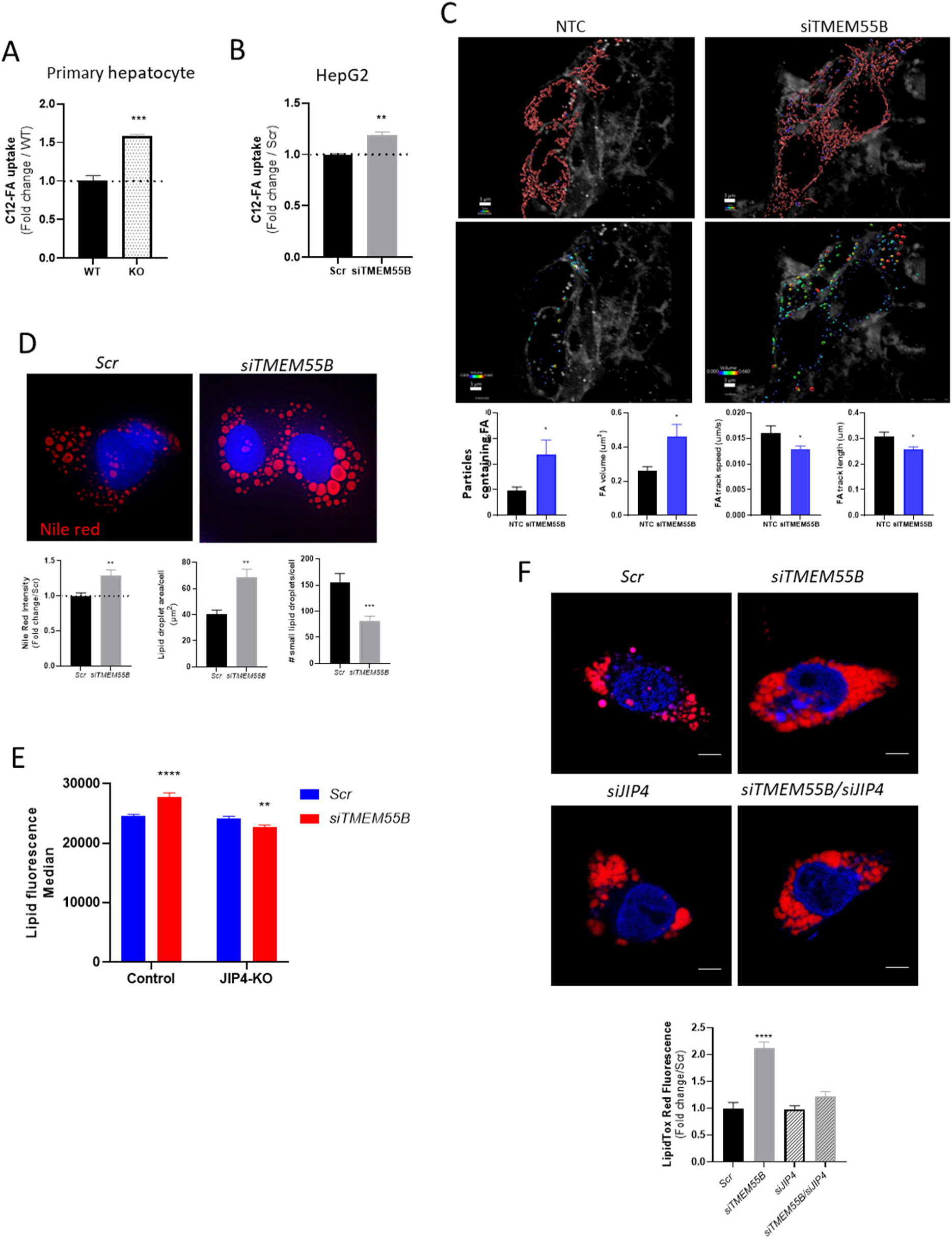
Tmem55b knockdown regulates cellular lipids through the lysosome position. A) Primary hepatocytes isolated from Tmem55b knockout (KO) and Tmem55b wildtype (WT) male mice, and B) HepG2 cells transfected with scr RNA or siRNA targeting TMEM55B were given 2mM BODIPY C12-FA for 30 mins and washed 3 times with PBS before FACS. C) After transfection HepG2 cells were incubated with 2mM BODIPY C12-FA for 30 mins before live cell imaging with Nikon spinning disk confocal microscope at 100X (Scale bars=3 μm) (Different colors in the images indicate different FA volume after surface reconstruction); D) Huh7 cells were transfected with TMEM55B and Scr siRNAs, incubated with 1mM oleate for 24hr and stained with Nile red (Scale bars=10 μm). E) Control and Jip4-knockout HepG2 cells were transfected with siRNAs targeting TMEM55B or scrambled controls, incubated with BODIPY493/503, and fluorescence quantified by FACS. Representative images are shown here. **F**) HepG2 cells were transfected with siRNAs targeting TMEM55B (siTMEM55B), JIP4 (siJIP4), siTMEM55B/siJIP4, or scrambled (Scr) siRNA. After 48 hr, cells were incubated with 1mM oleate for 24 hr, stained with LipidTOX Red, and imaged with Zeiss 710 confocal microscope at 63X (Scale bars=10 μm). Representative images are shown. Results are presented as mean ± s.e.m. N=3-12 per treatment group. *p<0.05, **p<0.01, ***p<0.001, ****p<0.0001 vs Scr by Student’s t-test.

We have shown that lack of *Tmem55b* led to hepatic steatosis *in vivo* (**Fig. 1A-C**). Based on the increased FA uptake and impaired mitochondrial fatty acid β-oxidation, we sought to confirm that loss of *TMEM55B* would also cause increased intracellular lipid accumulation *in vitro*. To test this, we incubated Huh7 cells with 1mM oleic acid and stained cells with Nile Red, a neutral lipid dye. After 24 hours there was 29% greater Nile red staining, 70% larger lipid droplet area per cell (p<0.01, **Supplemental Fig. S6B, Fig. 5D**), and fewer smaller lipid droplets with *TMEM55B* knockdown (p<0.001, **Fig. 5D**), which is consistent with prior reports that only smaller lipid droplets can be engulfed by lysosomes through lipophagy^53^. We confirmed that *TMEM55B* knockdown led to greater cellular steatosis in HepG2 cells incubated with BODIPY C12-FA (**Supplemental Fig. S6C, S6D**).

Given the known role of TMEM55B in lysosome motility^22,24,63^, we next tested whether the observed cellular steatosis was dependent on the ability of TMEM55B to impact lysosome movement. Under normal conditions (not stimulated with FA), *TMEM55B* knockdown decreased the speed, track length and duration of lysosomes in HepG2 cells (**Supplemental Fig. S3D**). TMEM55B enables lysosome transport by recruiting JIP4, a dynein scaffold, to the lysosome^22^. Thus, we created a HepG2 *JIP4* knockout (KO) cell line using CRISPR-Cas9, which led to an 86% reduction in *JIP4* transcript levels compared to HepG2 cells treated with a non-targeting control (NTC) gRNA (**Supplemental Fig. S6E**). While*TMEM55B* knockdown led to increased lipid accumulation in the NTC gRNA treated cells incubated with BODIPY C12-FA for 24 hours, knockdown had no effect on lipid accumulation in the *JIP4*-KO cell line (**Fig. 5E**). A similar lack of lipid accumulation was also seen after *TMEM55B/JIP4* siRNA-mediated double knockdown in HepG2 cells using flow cytometry (**Fig. 5F**, **Supplemental Fig. S6F**)

## DISCUSSION

In this study, we defined a new role for TMEM55B in mediating lipid metabolism through selective autophagy. We provide multiple lines of evidence, including two mouse models, primary mouse hepatocytes, and cultured human hepatoma cell lines, to support our findings. First, examining *Tmem55b* knockdown and knockout mouse models, we discovered that loss of hepatic TMEM55B led to excess hepatic lipid accumulation and greater progression to fibrosis.

These findings were consistent with our observations that iPSC-derived hepatocyte like cells from MASH patients had reduced *TMEM55B* transcript levels compared to healthy controls. Second, to investigate the mechanism by which TMEM55B deficiency accelerates MASLD onset and progression to MASH, we performed a series of *ex vivo* and *in vitro* studies which support the model that by preventing binding with the adaptor protein JIP4, responsible for facilitating lysosome movement, loss of TMEM55B increases lipophagy, leading to excess FA delivery to mitochondria and resulting in mitochondrial damage. This effect is exacerbated by impaired mitophagy, ultimately leading to disrupted mitochondrial networks, reduced FA oxidation, and increased oxidative stress. Together, these findings demonstrate how impaired lysosome trafficking can cause a cascade of metabolic defects via disruptions in lipophagy and mitophagy, resulting in the development of MASLD.

Lysosomes normally fuse with endosomes or autophagosomes in the perinuclear region^64^, after which lysosomal hydrolytic enzymes degrade the components to enable the release of materials^65^. We previously reported that in human hepatoma cell lines incubated with oleic acid, knock-down of TMEM55B led to the formation of enlarged lysosomes clustered around the nucleus. Here, we show that these enlarged lysosomes are filled with lipids, aka lipolysosomes. Lipolysosome formation could be attributed to impaired degradative activity within the lysosome, impaired release of lysosomal contents and/or increased delivery of lipids to the lysosome. We found no evidence of impaired lysosomal enzyme activity or lysosomal acid lipase activity (the enzyme that degrades triglycerides into free fatty acids^39^), or impaired release of FA from lysosomes. In fact, both lysosomal enzyme activity and FA release were increased. In contrast, there was greater co-localization and a higher number of direct contacts between lipid droplets and lysosomes, greater co-localization of lipids with LC3B (an autophagosome marker), and reduced p62 levels (a marker of autophagy). These findings indicate that loss of TMEM55B causes lipolysosomes formation through both macrolipophagy, an autophagosome mediated process^66^, and microlipophagy, an autophagosome-independent process of direct lipid extrusion from lipid droplets to lysosomes^67^. While hepatic steatosis in MASLD is typically characterized by excess accumulation of lipids within lipid droplets, MASLD patients have also been shown to have excess lipolysosomes^44,68–70^. Importantly, while lipolysosome formation is positively correlated with the MASLD activity score and fibrosis stage^70^, it was previously unknown how lipolysosomes formation relates to MASLD disease^70^ etiology. Our findings demonstrate how formation of lipolysosomes can be indicative of impaired lysosomal mobility, which ultimately impacts multiple aspects of cellular lipid metabolism, including increased lipophagy and impaired mitochondrial fatty acid oxidation, leading to the development of MASLD.

Typically decreased lipophagy is associated with MASLD onset. Impaired lipophagy has been observed in the livers of MASLD patients^68^, and pharmacological or genetic inhibition of lipophagy leads to hepatic steatosis and progression to MASH in animal models^47^. Induction of lipophagy in the liver can ameliorate MASLD through increased fatty acid oxidation^71^ and extracellular lipid secretion^49^, and thus has become a target for the development of novel MASLD theraputics^72,11^. While these long-standing observations appear to contradict our findings that inhibition of TMEM55B led to both increased lipophagy and greater development of MASLD, this discrepancy is likely attributed to the fact that TMEM55B impacts both lipophagy and mitochondrial function through mitophagy. Free fatty acids (FFA) are highly toxic to cells^73,74^, thus during lipophagy FA released in bulk are first esterified where they can be safely incorporated into lipid droplets^75^, and subsequently re-released by lipases prior to trafficking to the mitochondria^48^ for β-oxidation^76^. Thus, while elevating lipophagy typically reduces hepatic steatosis, this effect is dependent on efficient oxidation of FA by mitochondria, which not only produces energy, but also serves as a pathway to remove FA from the cell.

Based on the known effects of TMEM55B in lysosomal positioning and autophagic flux, we postulated that loss of TMEM55B would impair mitophagy and inhibit overall mitochondrial function. Consistent with this hypothesis, we found that loss of TMEM55B resulted in reduced mitochondrial volume, number, and membrane potential, fragmented mitochondrial networks, and impaired FA β-oxidation. There are many known regulators of mitophagy, including the autophagy adaptor protein p62/sequestosome-1. During mitophagy, p62 is recruited to damaged mitochondria, targeting them for lysosome decay^77^. Thus, lack of p62 has been shown to impair mitophagy^60,77^. We found that TMEM55B reduced levels of p62 and reduced lysosome-mitochondria co-localization, indicative of impaired mitophagy. In addition, loss of TMEM55B caused excess delivery of FA to mitochondria (in part due to elevated lipophagy). FA must be first converted to acylcarnitine in order to enter the mitochondria^78^, and acylcarnitine is toxic to mitochondria^56^, thus excess FA delivery to mitochondria has been shown to cause mitochondrial damage ^32^. Thus the dramatic effects of TMEM55B inhibition on mitochondrial health is due to the combined effect of excess lipophagy-created FFA causing mitochondria toxicity and the inability to remove and replace damaged mitochondria.

Hepatic mitochondrial dysfunction has been linked to the development and progression of MASLD^9,79^ as impaired mitochondrial FA oxidation and elevated oxidative stress has been observed in MASH patients and mouse models of MASLD^79–83^. While this dysfunction was historically thought to occur after steatosis, it has now been shown that mitochondrial damage can occur prior to the development of steatosis^84^. Increasing hepatic mitochondrial FA oxidative function in murine models of MASLD reversed hepatic steatosis and fibrosis^85–87^, demonstrating the importance of mitochondrial function in MASLD development. Mitochondrial dysfunction can contribute to MASH through the formation of ROS^88^, which can cause lipid peroxidation leading to the formation of aldehydes that stimulate mitochondrial depolarization^81^. Typically, this feed-forward loop is disrupted by mitophagy to remove and replace the damaged mitochondria. However, this process is disrupted in the absence of TMEM55B. ROS can also disrupt proteins and DNA, induce necrosis and apoptosis of hepatocytes, amplify the inflammatory response by activating Kupffer cells and recruiting circulating monocytes^89^, and importantly activate hepatic stellate cells responsible for promoting fibrosis^90^. Accumulation of defective mitochondria can also cause release of mitochondria-derived damage-associated molecular patterns (mito-DAMPs)^91^ or mtDNA^92^, inflammasome activation^93^, ER stress^94^, cell death and tissue damage^12^, all of which contribute to the observed relationship between mitochondrial dysfunction and MASLD onset and progression to advanced disease. Given the numerous mechanisms by which mitochondrial dysfunction promotes MASLD progression to MASH, it is not surprising that loss of TMEM55B, which enables accumulation of damaged mitochondria, also increases susceptibility to MASH.

Lastly, *Tmem55b* knockout has previously been reported to be lethal in mice during embryogenesis. Hashimoto Y. et al. replaced *Tmem55b* exons 2 to 6 with IRES-lacZ and PGK-neo-poly(A)-loxP cassettes in mouse embryonic stem cells^23^. We created our *Tmem55b* knockout mice by inserting loxP sites flanking *Tmem55b* exons 1 to 6 and crossed with Sox-2-cre mice to generate whole-body *Tmem55b^−/−^* (KO) mice. No Tmem55b protein was detected in the liver, adipose, or muscle of the Tmem55b^−/−^ mice, which are viable and fertile. In addition, recently, two paralogs of *Tmem55b* were knocked out in zebrafish, which were also viable and fertile^26^.

In summary, here we demonstrate that loss of *TMEM55B* led to MASLD onset and progression through enhanced lipophagy and impaired mitophagy. Notably, our findings here suggest that when mitochondria function is impaired, increased lipophagy promotes rather than ameliorates MASLD onset and progression. These findings illustrate the importance of maintaining mitochondrial function when targeting lipophagy as a MASLD therapeutic strategy.

## Acknowledgments

Sample processing was performed with the assistance of the UCSF Liver Center P30 DK026743, the UCSF MLK Cores Research Facility, and the UC Berkeley Electron Microscopy Core. Mouse RNAseq data was generated at QB3 Genomics, UC Berkeley, Berkeley, CA, RRID:SCR_022170. We thank Dr. Jacquelyn Maher for her input on iPSC-Hep RNAseq data analysis.

This work was supported by NIH R01 HL139902 (MM), R01 DK130391 (MM), R56 DK135259 (MM). NIH R01 EY030668 (AL), and National Eye Institute Diversity Supplement R01EY030668S1 (NLC). Additional funding to MM was provided by the Helen Fuller Springer Memorial Foundation through the UCSF Research Evaluation and Allocation Committee (REAC).

## Author contributions

Conceptualization: M.M. and Y.Q.

Methodology: Y.Q., N.L.C., N.Y.,

Formal analysis: Y.Q., A.M., E.T.

Investigation: Y.Q., S.T., N.L.C., N.Y., A.M., L.V., J.S.

Resources: A.L.

Writing – Original Draft: M.M and Y.Q.

Writing – Reviewing & Editing: all authors

Visualization: Y.Q., M.M.

Supervision: M.M., R.M.K, A.L.

Project Administration: M.M.

