## Supplemental data for "Loss of TMEM55B modulates lipid metabolism through dysregulated lipophagy and mitochondrial function"

Supplementary Figure S1

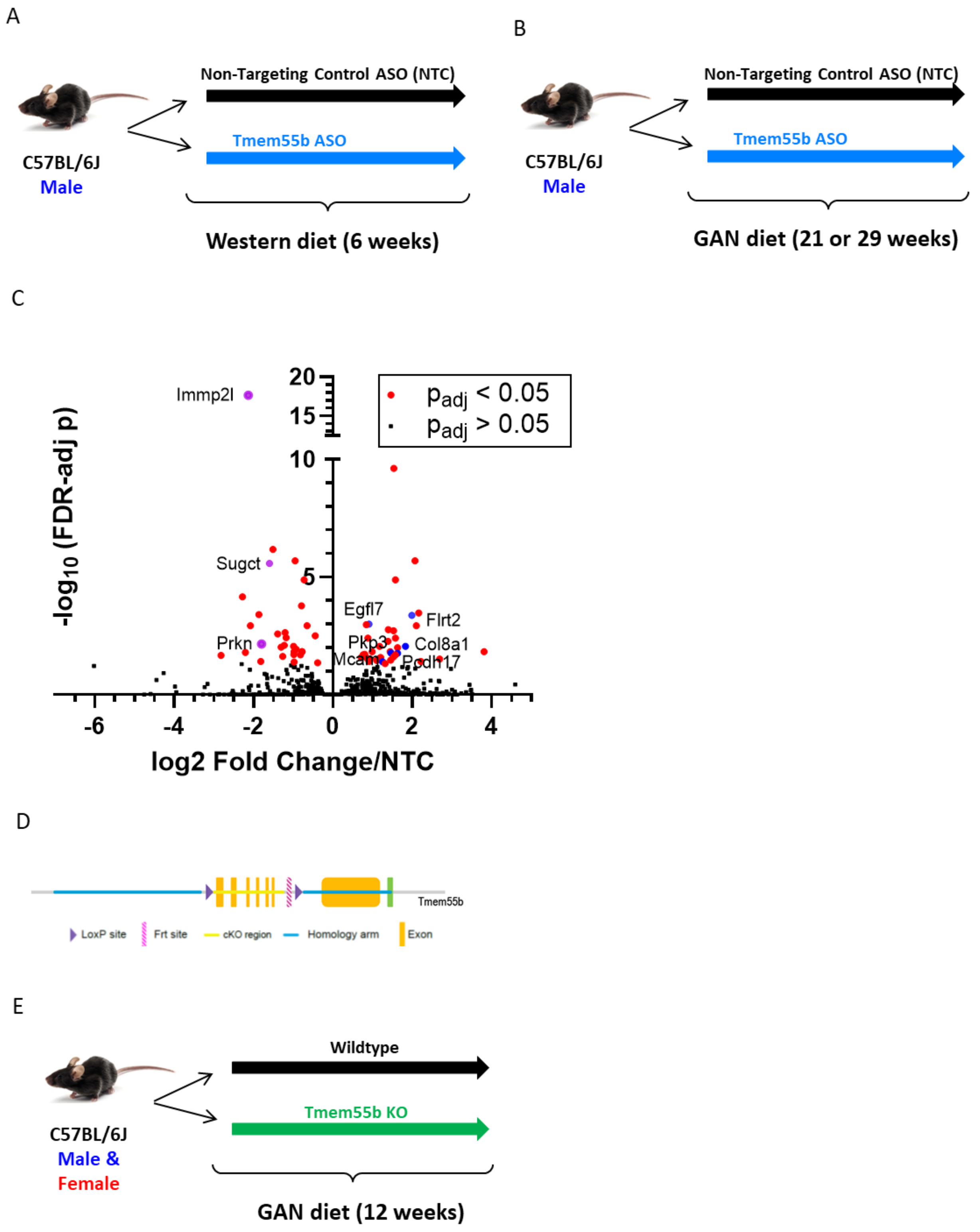

**Supplementary Fig. S1.** Study design with different mouse models and Liver RNAseq results. A) Study design of in vivo Tmem55b knockdown. Six-week old male C57BL/6J mice were treated with an ASO against Tmem55b (Tmem55b-ASO) or a non-targeting control (NTC) at a dose of 25 mg/kg body weight/week and fed a Western diet (0.2% cholesterol, 42% fat) for 6 weeks. B) Male C57BL/6J mice were treated with Tmem55b-ASO or NTC-ASO and fed Western diet for 21 or 29weeks. C) RNAseq results from the liver of Tmem55b-ASO and non-target ASO treated male mice. Six-week old male C57BL/6J mice were treated with an ASO against Tmem55b (Tmem55b-ASO) or a non-targeting control (NTC) at a dose of 25 mg/kg body weight/week and fed a Western diet (0.2% cholesterol, 42% fat) for 4 weeks. Total RNA was extracted from mouse livers (n=4/group), checked for quality on a Bioanalyzer, and made into polyA-selected, strand-specific RNA-seq libraries for 150 bp paired-end sequencing on Illumina NovaSeq machines. DESeq2 was used for differential expression analysis, and p-values were adjusted for multiple testing using a false discovery rate (FDR) approach. 68 of the 17,903 genes tested were significantly differentially expressed at an FDR=5% (38 upregulated and 31 downregulated. Purple color dots are the genes (Prkn, Immp2i, and Sugct ) which regulate mitochondrial function, while blue color dots are upregulated genes involved in cell adhesion (*Flrt2*, *Egfl7*, *Col8a1*, *Pkp3*, *Pcdh17, Mcam*). D) Creation of Tmem55b knockout mice and the effect of TMEM55B knockdown in lipophagy. D) Schematic of the Tmem55b knockout allele. We created a *TMEM55B* knockout model by inserting loxP sites flanking Tmem55b exons 1 to 6 in a C57BL/6N background. The Tmem55b$^{fl/+}$ mice were backcrossed 7 times to the C57BL/6J strain, and subsequently crossed with Sox-2 mice to generate whole-body Tmem55b$^{-/-}$ (KO) mice on a >99% C57BL/6J background. E) Six-week-old male and female wildtype and TMEM55B knockout mice were put on GAN diet for 12 weeks.

### Supplementary Figure S2

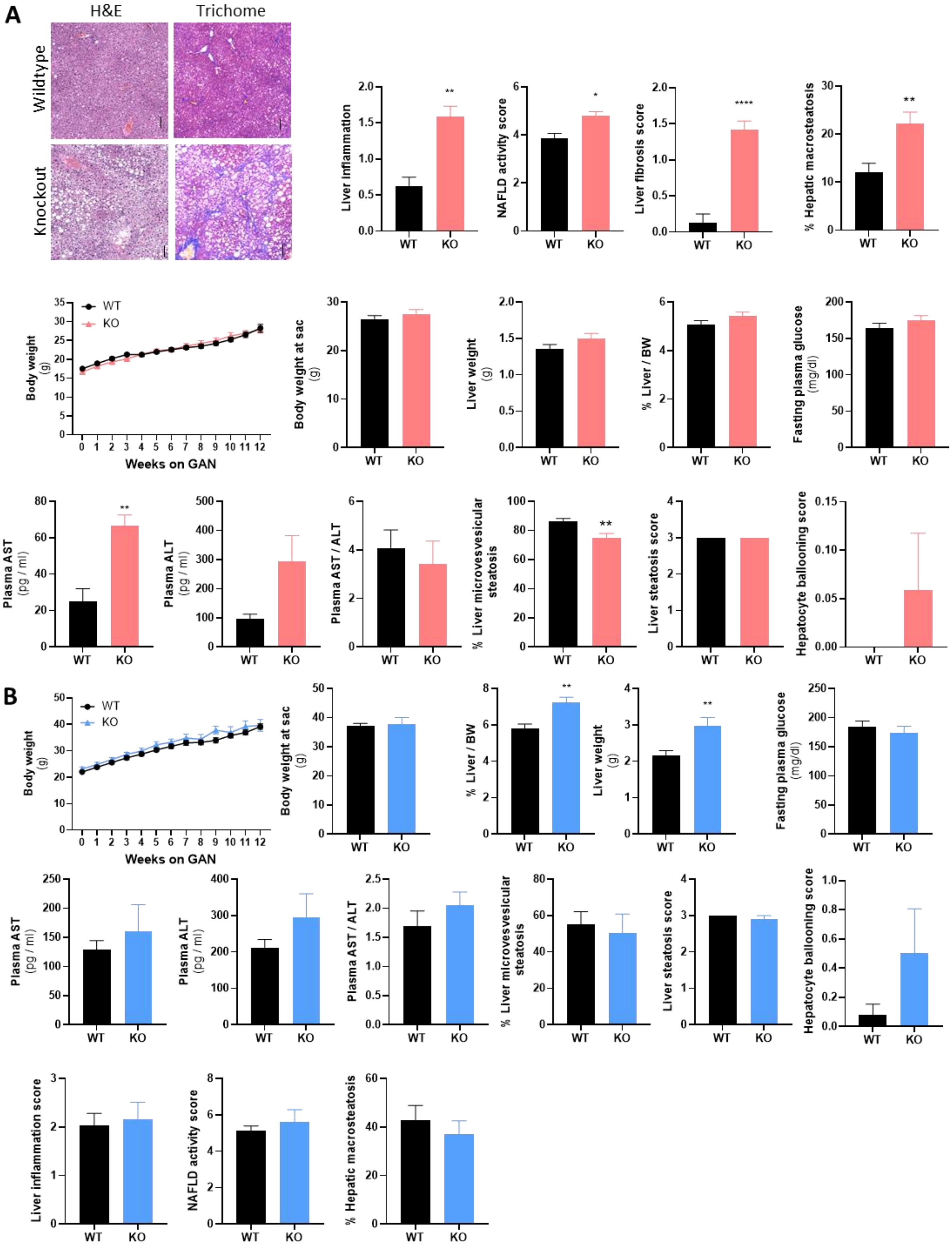

**Supplementary Fig. S2.** Phenotypes of female and male WT and Tmem55b KO mice. Mice were fed a GAN diet starting at 6-week-old for 12-weeks. Live histology of female (A) and male (B) mice was evaluated by a blinded pathologist. Scale bar=100 µm. Body weight, liver weight and fasting glucose were measured in both female (A) and male mice (B). Plasma AST, ALT, and AST/ALT were detected by Liasys 330 Clinical Chemistry Analyzer in female (A) and male (B) mice. Live histology of female (A) and male (B) mice was evaluated by a blinded pathologist. Scale bar=100 µm. N=9-13. For the animal studies, quantified results are presented as mean ± s.e.m. **p<0.01 vs. Tmem55b+/+ by Student's t-test.

**Supplementary Figure S3**

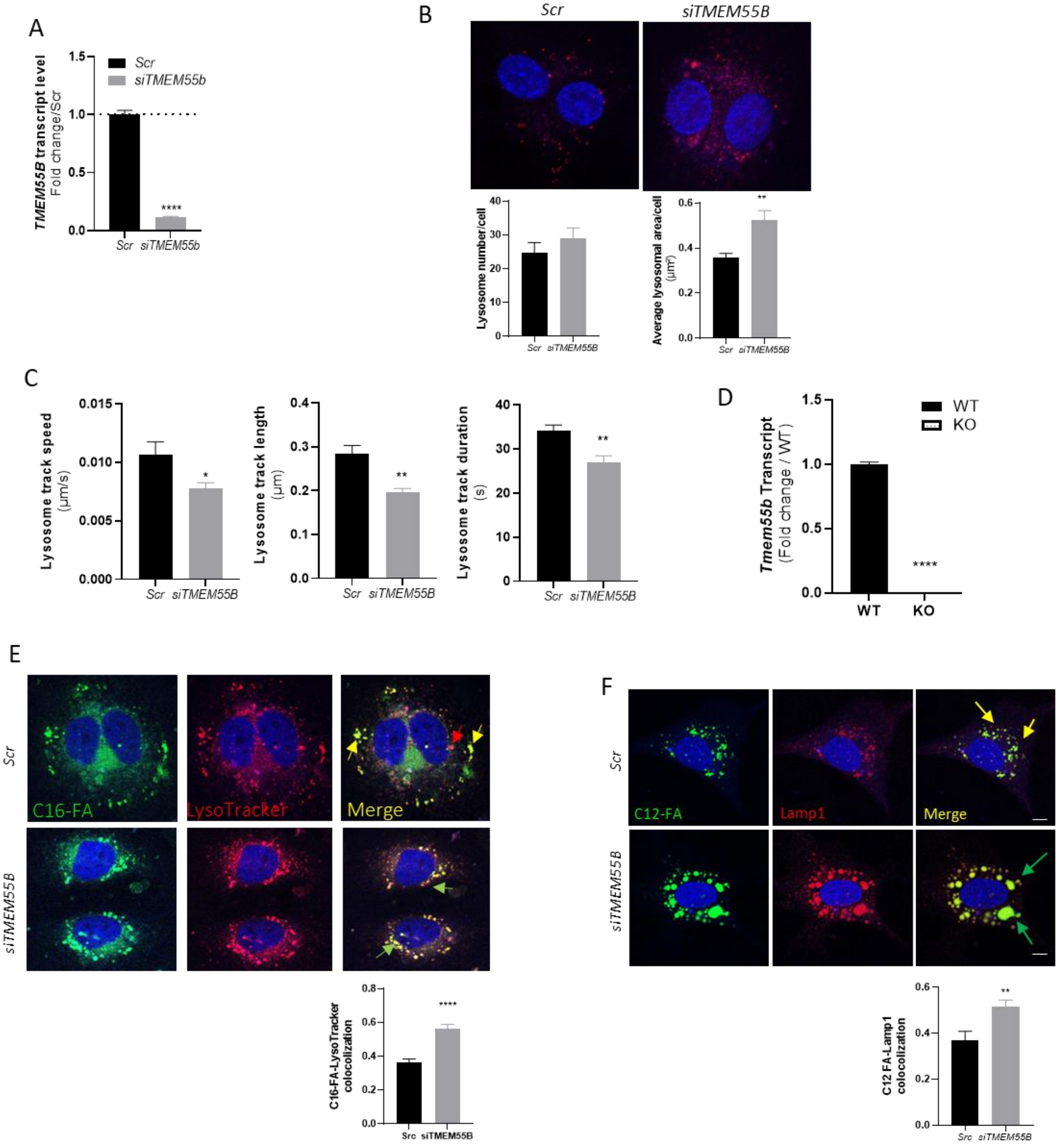

**Supplementary Fig. S3.** The effects of TMEM55B knockdown on lysosomes and lipid accumulation inside lysosomes. HepG2 cells were transfected with TMEM55B or Scr siRNAs. After 48 hr, TMEM55B transcript levels were quantified with qPCR (A), or cells were statined with LysoTracker for quantification of lysosome number and areas per cell (B). (C) Lysosome speed, track length and duration was measured with 1 mM palmitate treatment for 1 hr after TMEM55B knockdown using live cell imaging. (E) TMEM55B transcript levels were quantified in Huh7 cells with qPCR 48 hr after siRNA transfection. After transfection,  HepG2 cells were given 2 µm of green BODIPY™ FL C16-FA (E) or BODIPY™ FL C12-FA (F) overnight, stained with LysoTracker (E) or anti-LAMP1 antibody (F) for 1 hr at room temperature followed by 2$^{nd}$ Goat anti-rabbit IgG (H+L) Alexa Fluor 568 for 1 hr, and examined by confocal microscopy. Representative images are shown here. Results are presented as mean ± s.e.m. *p<0.05 **p<0.01, ****p<0.0001 vs Scr by Student's t-test.

**Supplementary Figure S4**

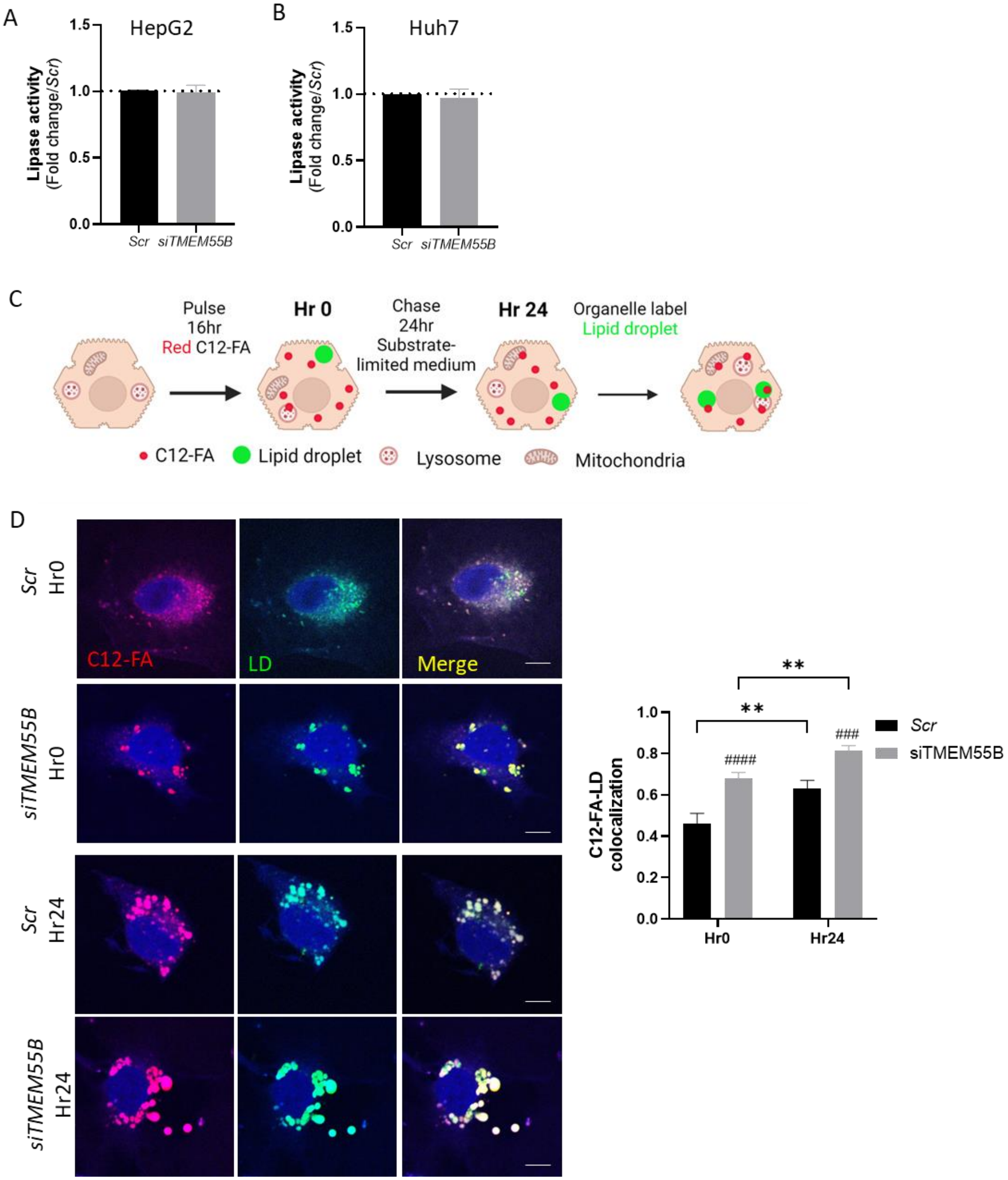

**Supplementary Fig. S4.** TMEM55B knockdown showed no effect on cytosolic lipase activity and increased FA trafficking to lipid droplets. After transfection, cellular lipase activities were measured in both HepG2 (A) and Huh 7 (B) cells using the Lipase Assay Kit from Abcam. C) Schematic representation of the FA pulse-chase assays. HepG2 cells were transfected with TMEM55B and Scr siRNAs, pulsed with Red C12-FA (1μM) overnight, washed, and chased with substrate-limited DMEM supplemented with 0.5 mM of glucose, 1 mM of glutamine, 0.5 mM of carnitine, and 1% FBS for 24 hours. After which, cells were labeled with 1 μM BODIPY 493/503 for lipid droplets, and imaged with Zeiss 710 confocal microscope (D). C12-FA localization to lipid droplets (D) before and after the chase were quantified. Representative images are shown here. Results are presented as mean ± s.e.m. **p<0.01 vs Hr0; ####p<0.0001 vs Scr by two-way ANOVA.

**Supplementary Figure S5**

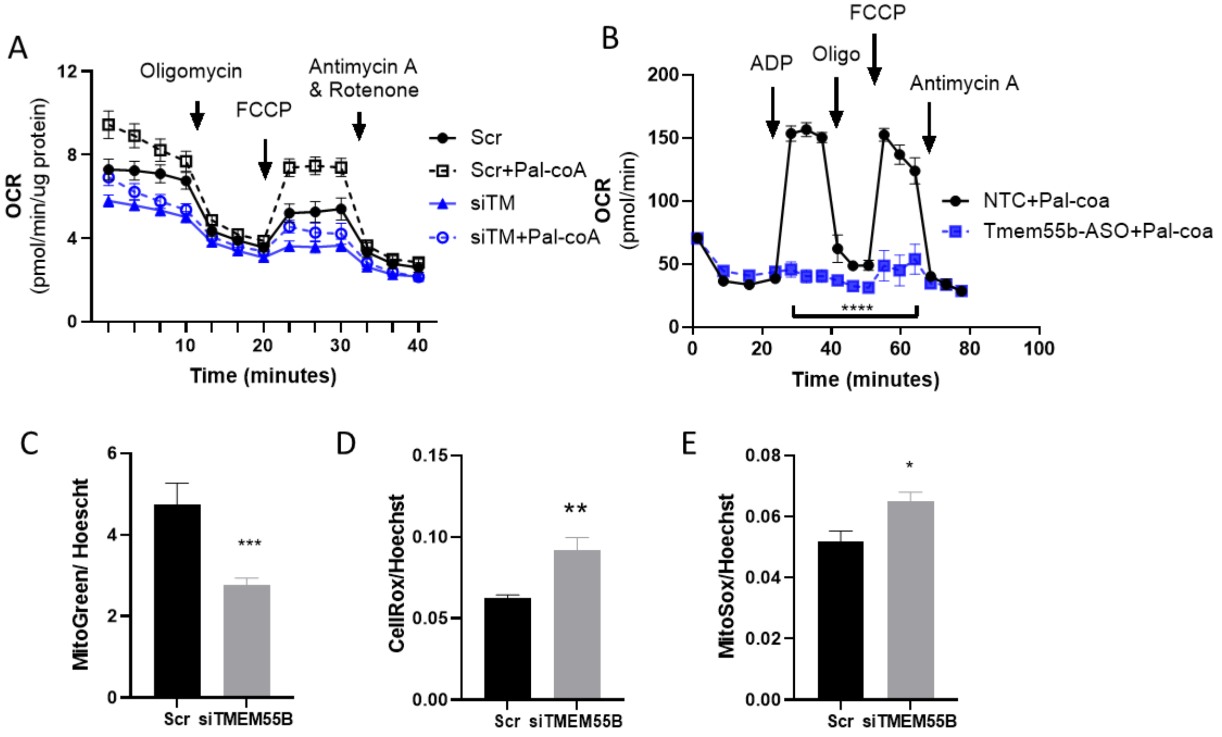


**Supplementary Fig. S5.** The effects of TMEM55B deficiency on mitochondria. A) HepG2 cells were transfected with siRNAs targeting TMEM55B (siTMEM55B) or scrambled (Scr) siRNA. After 48 hr, cells were treated with XF Plasma Membrane Permeabilizer (PMP) and oxygen consumption rate (OCR) was measured by Seahorse XF Analyzer with and without with or without the addition of 40 µm palmitate-coA. B) C57BL/6J male mice were and fed a GAN diet (40% kcal fat, 20% kcal fructose, 2% cholesterol) for 21 weeks since 6-week-old. The animals were i.p. injected with 25 mg/kg body weight/week of antisense oligonucleotides (ASO) targeting *Tmem55b* or a non-targeting control (Ionis Pharmaceuticals). Mitochondria were isolated from the liver tissues of 27-week-old male mice, OCR was measured by Seahorse XF Analyzer. After transfection, HepG2 cells were treated with 200nM MitoGreen for 30 min (C), 5 µM CellRox Green for 30 min (D), or 5 µM MitoSox Red for 10 min (E), and fluorescence levels were quantified with BioTek Synergy H1 Plate Reader. Results are presented as mean ± s.e.m. **p<0.01, ***p<0.001, ****p<0.0001 vs Scr or NTC by Student's t-test or One-way ANOVA.

**Supplementary Figure S6**

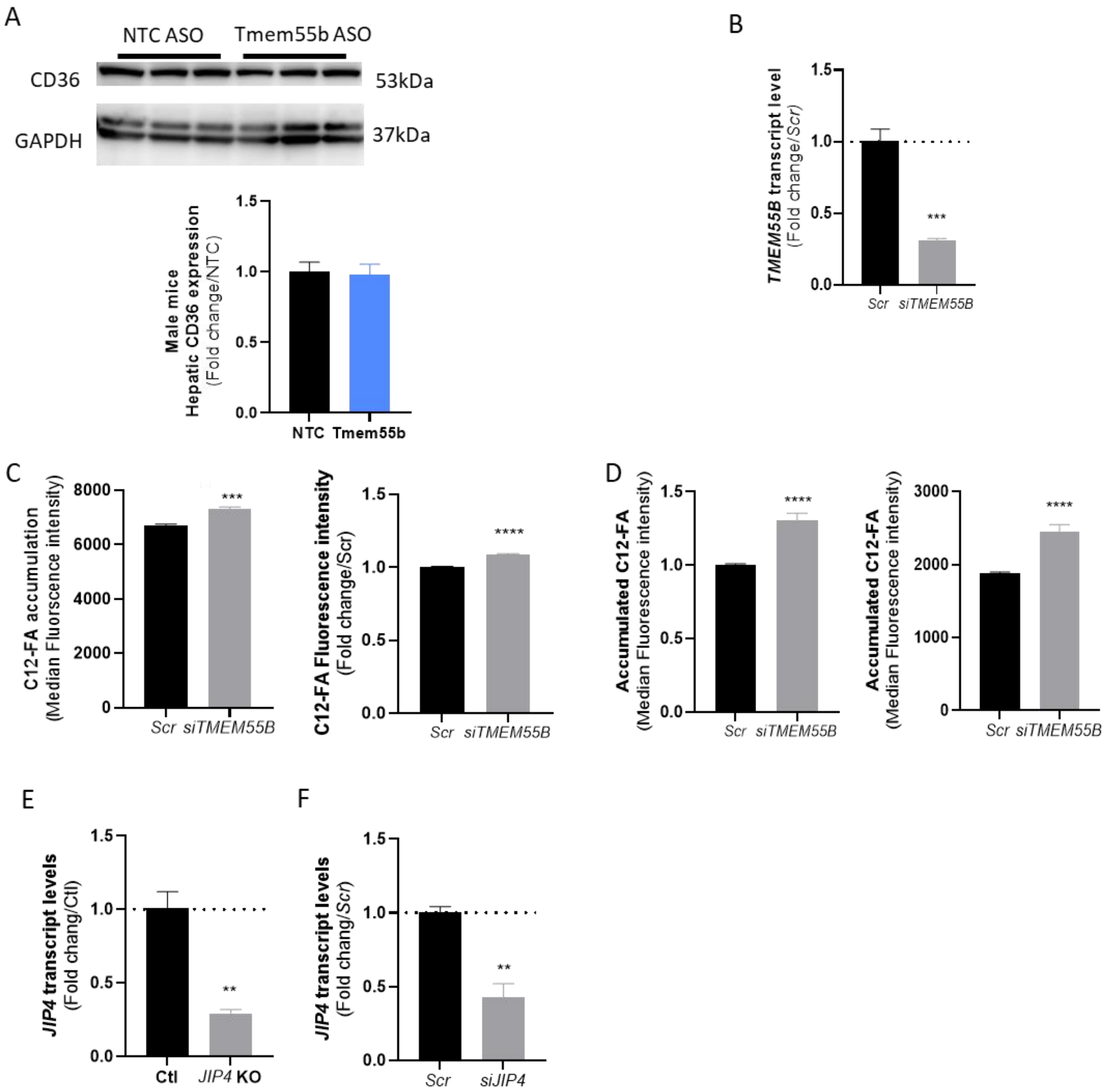

**Supplementary Fig. S6.** TMEM55B knockdown led to increased FA uptake and accumulation. (A) Protein levels of CD36 in the livers between ASO-Tmem55b and ASO-NTC treated male by Western blot. (B) Huh7 cells were transfected with siRNAs targeting TMEM55B (siTMEM55B) or scrambled (Scr) siRNA. After 48 hr, TMEM55B transcript levels were measured by qPCR. After transfection, HepG2 cells were incubated with 2μM Green BODIPY C12-FA for16 hr (C) and chased with substrate-limited medium for 24 hr (D) before fluorescence quantified by FACS. JIP4 transcript levels were quantified by qPCR in HepG2 control (Ctl) and JIP4 knockout (KO) cell line(E) and scrambled siRNA (Scr) and siRNA targeting JIP4 (siJIP4) cells (F) . Results are presented as mean ± s.e.m. $^{**}p<0.01$, $^{***}p<0.001$, $^{****}p<0.0001$ vs Scr or NTC by Student's t-test or One-way ANOVA.

**Supplementary Figure S7**

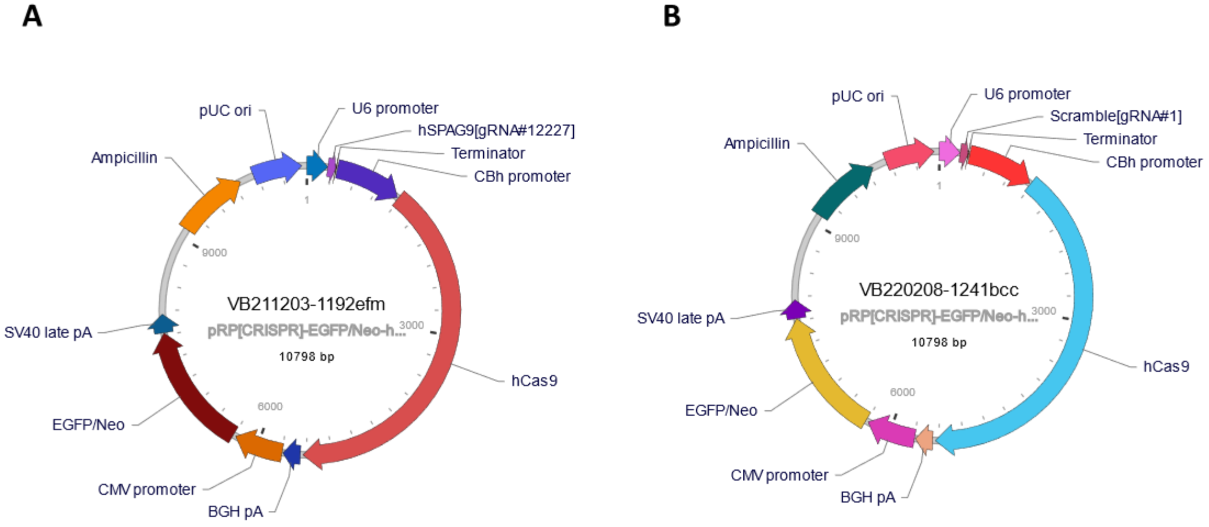


**Supplementary Fig. S7.** The design of single gRNA targeting JIP4 (VB211203-1192efm, A) and non-targeting gRNA as negative control (VB220208-1241bcc, B) from VectorBuilder.
